# Culture Dimensionality Governs Gallium Maltolate Response in Glioblastoma: Comparative Analyses in 2D and 3D Models

**DOI:** 10.1101/2025.09.17.676610

**Authors:** Paulina Szeliska, Karol Jaroch, Weronika Wróblewska, Łukasz Kaźmierski, Małgorzata Maj, Barbara Bojko

## Abstract

Gallium maltolate (GaM) targets iron-dependent processes in glioblastoma (GBM), but responses vary with model context. We evaluated GaM across established (A-172, U-87 MG) and patient-derived (3005, 3019, 3034, 3048, 3073) GBM lines in 2D and 3D using viability modelling (IC10/IC50/IC90), TFRC quantification, oxygen consumption rate (OCR), and PCA/PLS-DA– guided metabolomics with FDR and VIP-based selection. GaM reduced viability in all models, with a right-shift of dose–response in 3D. TFRC correlated with IC50 in 2D but not 3D, unless excluding U-87 MG from the analysis. OCR was markedly suppressed in A-172, U-87 MG, 3048, and 3073, particularly in 3D, while 3005 and 3019 were more respiration-resilient. Multivariate analyses showed treatment-dominant separation in 3005/3048, format dominance in 3019/3034, and time effects in A-172/U-87 MG/3073. A concise metabolic signature (tryptophan, methionine, uracil, allantoin) indicated coordinated perturbations in amino acid, nucleotide, and redox pathways. These findings support 3D, patient-derived systems for more predictive GaM evaluation.

## 1. Introduction

Glioblastoma (GBM) is a very heterogeneous tumor associated with a poor survival rate. According to WHO’s classification for 2021, histological analysis of tissue is no longer sufficient to classify a central nervous system tumor^1^. GBM diagnosis includes investigation of mutation in isocitrate dehydrogenase (IDH-wildtype) and three genetic characteristics: TERT promoter mutation, EGFR amplification, and +7/-10 chromosome copy number. This new grading system allowed classifying histologically II or III grade diffuse gliomas into IV grade. Standard GBM treatment is composed of tumor resection followed by radiotherapy and temozolomide chemotherapy. However, research on molecular subtypes of glioblastoma suggested further differentiating them based on molecular changes. It would allow for personalized treatment. Novel GBM therapy focuses on a targeted approach. Immunotherapy, molecular-targeted, and angiogenesis-targeted approaches are most intensely researched^2^. However, bevacizumab is the only targeted drug for recurring GBM treatment approved so far. There is still a need for new therapeutic approaches to improve the survival rate of patients^3^.

Gallium was first introduced in the medical field in the late 1960s as a radioisotope^4^. However, some Ga(III) complexes showed potential in oncological therapy. Ga(III) shares some properties with iron and can be introduced into metabolic pathways as an iron substitute. Ga(III) can bind to iron-binding proteins and cause iron deficiency in rapidly proliferating cells such as cancer cells^5^. Transferrin (Tf) is a protein with two iron-binding sites and is responsible for delivering iron to cells; therefore, it is a potential way to provide Ga-based compounds as well. Approximately one-third of the circulating Tf is an iron-loaded Tf, also known as holo-Tf. This loaded Tf enters the cell by binding to the Tf receptor (TfR1 and TfR2), and the non-loaded Tf (apo-Tf) remains circulating in the organism. It leaves about 2/3 of Tf available to deliver Ga(III)-complexes into cells ^6,7^. Nowadays, researchers focus on three Ga-complexes with the highest potential in clinical applications, and they are currently undergoing clinical trials: gallium nitrate (GaN), tris(8- 8- quinolinolato)gallium(III) (KP46), and gallium maltoalte (GaM) ^8^. GaM is a complex of a gallium ion and three deprotonated gallium groups. It was proven to have a few times higher bioavailability than gallium salts^9^. It is an orally administered compound due to its high bioavailability; moreover, most of it is Tf-bound in the blood^10^. It has been proven to inhibit cell proliferation in lymphoma resistant to GaN ^11^. Lately, Chitambar et al. analyzed the mechanism of action of GaM in 2D and 3D glioblastoma cell culture, as well as in tumor rat xenograft, and showed the potential of GaM to induce tumor cell apoptosis via disrupting the iron homeostasis. The ability of gallium to cross the blood-brain barrier (BBB) by binding to endogenous Tf has enhanced delivery and targets TfR-bearing GBM. GaM has been shown to influence mitochondrial function of tumor cells as well as RRM2 activity, leading to blocking DNA synthesis in GMB cells without affecting normal cells. GaM has an impact on the TCA cycle and inhibits mitochondrial oxygen consumption^12^. A phase 1 clinical trial is being conducted to determine the response of patients with relapsed and/or treatment-refractory tumors ^13^.

Metabolic alterations in cancer cells have long been explored for their usefulness in profiling the phenotypes of many tumors. Research revealed a good correlation between mutations found in GBM, e.g., PDGFRA, IDH1, EGFR, and NF1 and the tumor’s metabolic fingerprint. Thanks to extensive work on determining possible GMB metabolomic profiles, it became a promising tool for preclinical drug screening and tumor resistance to therapy exploration. However, the correlations between tumor metabolomic profile, TFRC, and the GaM treatment response have not yet been made. The comparison of the metabolomic profile with the results of standard cell culture assays will help to evaluate potential onco-metabolic targets for assessing the efficiency of the GaM therapy.

SPME is not a commonly used sample preparation method, yet it is an extremely promising tool for in vitro pharmaceutical research, as proven in previous studies conducted in our laboratory ^14,15^ and other studies^16,17^. Moreover, SPME was previously used for the analysis of brain and brain tumors in vivo^18–23^. Results revealed that SPME can provide spatially resolved metabolic and lipidomic profiles of patients’ brains in vivo by utilizing minimally invasive sampling, which omits physical sample consumption. Due to this unique feature, the technique is also known as “chemical biopsy”. In addition, the use of specially optimized biocompatible coating enables covering both hydrophobic (e.g., lipids) and polar compounds (e.g., amino acids)^18^. The same method was proposed for the studies of brain tumors. SPME in fiber form was used to penetrate tumor tissue for metabolome/lipidome sampling. The results showed that this sample preparation method was capable of providing a characteristic phenotypic snapshot of the brain tumor, providing lipidomic^22^ and metabolomic^23^ markers of disease. Following these investigations, targeted lipidomic analysis with the use of Coated Blade Spray was carried out. The results showed different levels of carnitine and acylcarnitines correlating with IDH-mutation and 1p/19q co-deletion status. Moreover, utilizing CBS, which omits LC separation and allows for fast instrumental analysis without compromising sample cleanup, demonstrated the potential of the approach for rapid on-site screening of potential biomarkers.^19–21^

Combining pharmaco-metabolomics with cell phenotype and genetic analyses in 3D culture models may advance understanding of glioblastoma, particularly within the iron-imbalanced microenvironment. Linking these profiles with therapeutic response could provide novel insight into gallium maltolate as a potential treatment. Applying SPME for the studies, which has proven its usefulness in glioma metabo-lipidomic profiling, *in vitro* and *in vivo* temporal studies, would help bridge the presented *in vitro* data with subsequent *in vivo* animal research.

## 2. Materials and Methods

Unless stated otherwise, all chemicals were purchased from Merck Group (Darmstadt, Germany).

### 2.1. Gallium malotlate (GaM) preparation

Gallium maltolate was purchased from THE BioTech (Los Angeles, CA, USA) and kept at 4°C. Stock solution of 1 mM was prepared by suspending in sterile, ultra-pure water, vortexing and 5 min sonication, followed by another thorough vortexing.

### 2.2. 2D Cell culture

Cell line U-87MG was purchased from American Type Culture Collection (ATCC, Manassas, USA), A-172 cell line was purchased from Cell Line Service (CLS, Eppelheim, Germany). Both cell lines were cultivated in DMEM (Corning, NY, USA) supplemented with 10% Fetal Bovine Serum (Corning, NY, USA) and Antibiotic Antimycotic Solution. Cells were cultured at 37°C, 5% CO2, and constant humidity. Patient-derived glioblastoma cells were acquired from the Human Glioblastoma Cell Culture resource (www.hgcc.se) at the Department of Immunology, Genetics and Pathology, Uppsala University, Uppsala, Sweden ^24^. The 3005, 3019, 3034, 3048, and 3073 cells (basic characetristics in Tab. S1) were cultured according to the HGCC guidelines; DMEM: F12 (high glucose):Neurobasal, 1:1 (Thermo Fisher Scientific Inc., Waltham, MA, USA) supplemented with N1 supplement (0.5x) and N2 supplement (0.5x), B27 supplement (1x), Epidermal Growth Factor (EGF, 10 ng/ml) and basic Fibroblast Growth Factor (bFGF, 10 ng/ml) and Antibiotic Antimycotic Solution. HGCC cells were cultured on culture dishes coated with polyornithine (10 µg/ml) and laminin (10 µg/ml). Cells were passaged at 70-80% confluence. U-87MG and A-172 cell lines were detached with trypsin, while accutase was used for HGCC cells. Cell counting was performed by mixing with trypan blue and counted by an automated cell counter (Countess® II FL, Invitrogen by Thermo Fisher Scientific Inc., Waltham, MA, USA).

### 2.3. 3D cell culture

3D cell cultures were established from the 2D cell cultures described above. The protocol described by Wanigasekara et al. was followed with minor changes^25^. Briefly, cells were counted with an automatic cell counter, and a cell suspension of 10000 cells/ 200µl was prepared, and 200 µl of cell suspension was added to a 96-well ultra-low attachment culture plate (Nunc™, Thermo Fisher Scientific Inc., Waltham, MA, USA). The plate was then spun (230 xg, 5 min) and cell clusters incubated for 24h, and then 100 µl of media was removed and an equal volume of fresh medium was added. 100 µl of cell culture media was changed every 2^nd^-3^rd^ day. Spheroids were cultured for 10 days before GaM was added.

### 2.4. Viability assay

Cell viability was assessed using the MTT assay (2D cultures) or the CellTiter-Glo® 3D assay (spheroid cultures). Cells of U-87MG, A-172, 3073, 3048, 3034, 3019, 3005 were seeded at a density of 2500, 2500, 7500, 10000, 10000, 7500, and 12000 cells/well, respectively, in a 2D culture and 10000 cells/well for spheroids (12000 for 3005 cell line). Cells were then incubated (24h for 2D and 10 days for 3D spheroids). After incubation, 100 µl of fresh cell culture media was added with GaM in a 15 to 165 µM concentration range. For 2D cells, the MTT assay was performed after 24h and 72h. Celltiter Glo was performed after 72h and 7-day incubation for spheroids.

For 2D cell culture, after incubation, the medium was changed for 100 μL of MTT solution (1 mg/ml in DMEM with no phenol red, 10% FBS) and incubated for 3h. Formed formazan crystals were dissolved in isopropanol (100 µl), and absorbance (570nm/690nm) was measured using a microplate reader (Synergy H1, BioTek, Winoosky, VT, USA). Cell viability was expressed as a percentage relative to the untreated control.

Cell viability in 3D spheroid cultures was determined using CellTiter-Glo® 3D Cell Viability Assay (Promega). Following treatment with GaM, spheroids were gently transferred with a cut pipette tip to a white 96-well plate. An equal volume of CellTiter-Glo® 3D reagent was added directly to each well. Plates were incubated for 30 min at room temperature on an orbital shaker to allow complete cell lysis and ATP stabilization. Luminescence, proportional to the amount of metabolically active cells, was measured using a microplate reader. Cell viability was expressed as a percentage relative to the untreated control.

### 2.5. Oxygen consumption assay

Extracellular oxygen consumption was measured using the Extracellular Oxygen Consumption Assay (OCR) Kit (Abcam, ab197243, Cambridge, UK) following the manufacturer’s protocol. Briefly, cells were seeded in black, clear-bottom 96-well plates at a density of 1×10^5^ cells per well and cultured overnight. After adding GaM, 10 µL of Extracellular Oxygen Consumption Reagent was added to each well. Plates were immediately sealed with the supplied mineral oil overlay to prevent oxygen diffusion and incubated at 37 °C. Fluorescence (Ex/Em = 380/650 nm) was measured kinetically using a microplate reader, and oxygen consumption rates were calculated from the change in signal over time.

### 2.6. Protein Extraction and Quantification

Cells were collected and transferred to 15 mL conical tubes. After centrifugation at 300 × g for 7 min at room temperature, cell pellets were resuspended in ice-cold PBS and centrifuged again at 300 × g for 7 min at 4 °C. Pellets were lysed in Complete Cell Extraction Buffer supplemented with protease inhibitors (1 mL per 1 × 10^8 cells). Lysates were vortexed briefly, incubated on ice for 30 min with intermittent mixing, and clarified by centrifugation at 13,000 × g for 10 min at 4°C. Supernatants were transferred to fresh tubes and stored at –80 °C. Total protein concentration was determined using the BCA Protein Assay Kit (Thermo Fisher Scientific) according to the manufacturer’s protocol.

### 2.7. Human Transferrin Receptor (TfR/CD71) ELISA

Following the manufacturer’s protocol, TfR/CD71 levels were quantified using the Human Transferrin Receptor ELISA Kit (Thermo Fisher Scientific Inc., Waltham, MA, USA). Based on BCA quantification, equal amounts of protein were diluted in assay buffer and loaded in duplicate into antibody-coated 96-well plates along with standards. After incubation at room temperature for 2.5h, wells were washed and incubated sequentially with biotinylated detection antibody and HRP-conjugated streptavidin. Signal was developed with TMB substrate, and reactions were stopped with 2 NH₂SO₄. Absorbance was measured at 450 nm using a microplate reader. TfR concentrations were calculated from the standard curve and normalized to total protein.

### 2.8. Cell lysis for endometabolome analysis

2D cells were collected during a standard passage procedure and counted by an automatic cell counter as described in 2.2. Cells were resuspended in PBS (1 x 10^6^ cells/ml). 3D spheroids were transferred into Eppendorf tubes and centrifuged, then resuspended in PBS (1 spheroid/100µl). Both 2D and 3D cells were submerged in liquid nitrogen for metabolome quenching and kept at - 80 °C until further analysis. Defrosted samples were sonicated (1 cycle, 5 min, 70%) and then cooled on ice (5 min). Extracts were centrifuged at 10,000 x g for 10 min, and the supernatant was transferred into fresh collection tubes. Extracts were then spiked with an internal standard L-Tryptophan-d8 (TRC, Vaughan, Canada) to monitor extraction in the final concentration of 100 ppb.

### 2.9. SPME-LC-MS/MS

Solid-phase microextraction (SPME) coated blade spray (CBS) devices (CB-HLB, 10 mm coating, Restek Cat No. 23248) were purchased from Anchem (Warsaw, Poland). Before use, the blades were preconditioned overnight in methanol: water (1:1, v/v) under static conditions. Immediately before extraction, CBS blades were rinsed for 5 s with ultrapure water. For extraction, 120 µL of spiked sample was agitated for 2 h at 850 rpm using a BenchMixer™ XLQ QuEChERs Shaker/Vortexer (Merck Group). After extraction, blades were briefly rinsed again in ultrapure water (5 s) before desorption. Analyte desorption was performed in acetonitrile: water (1:1, v/v) for 2h under agitation at 850 rpm. Desorbed samples were analyzed using a Nexera UHPLC system (Shimadzu, Kyoto, Japan) coupled to an LC–MS 8060 triple quadrupole mass spectrometer (Shimadzu, Kyoto, Japan). The LC–MS/MS method was based on the Primary Metabolites Version 3 method package (Shimadzu, Kyoto, Japan). Chromatographic separation was performed on a reversed-phase Discovery HS F5 column (100 mm × 2.1 mm, 3 µm; Supelco, Bellefonte, PA, USA). Quality control (QC) was ensured by including pooled QC samples and probe blanks throughout the study

### 2.10. Data Processing and Statistical Analysis

Dose–response curves for GaM were generated from MTT and CellTiter-Glo viability assays using R (tidyverse, drc, and stringr packages). Inhibitory concentrations (IC10, IC50, IC90) were estimated using the drc package ED function.

Transferrin receptor (TFRC) expression levels were quantified, and statistical significance of group differences was assessed in R using tidyverse, ggplot2, ggpubr, dplyr, and FSA packages. Comparisons between groups were performed with a two-tailed Student’s *t*-test, with *p* < 0.05 considered statistically significant.

The association between baseline TFRC levels in untreated control cells and GaM sensitivity (IC10 and IC50 values) was evaluated using Pearson’s correlation test, implemented in R with tidyverse and ggpubr packages.

LC-MS/MS raw data were processed with Skyline software with MRM transitions provided in the Primary Metabolites Version 3 method package (Shimadzu, Kyoto, Japan).

Pooled QC and blank samples were used for data pre-processing. Pre-processed data sets were implemented into Metaboanalyst 6.0, a free online software^26^. Metaboanalyst’s batch correction module was used in automated mode. Batch corrected data were further normalized based on internal standard and analyzed in Metaboanalyst, Statistical Analysis [one factor] with the following parameters for normalization node: sample normalization was set to none, data were log10 transformed, and auto scaling data scaling was implemented. Principal component analysis (PCA) and partial least squares discriminant analysis (PLS-DA) were performed to evaluate the separation among sample groups. Variables with variable importance in projection (VIP) scores exceeding 1 were designated as significant, reflecting their contribution to the model’s discriminative power and predictive reliability.

Further statistical analyses were performed in R (version 4.5.0) using the dplyr, ggplot2, ggpubr, ggsignif, tibble, tidyverse, FSA, and purrr packages. The Kruskal–Wallis test followed by Dunn’s post hoc test with false discovery rate (FDR) correction was applied for four-group comparisons. For two-group comparisons, the Wilcoxon rank-sum test with FDR correction was used. Metabolites were considered significant if they passed the statistical threshold and exhibited a variable importance in projection (VIP) score greater than 1.

## 3. Results

### 3.1. TFRC Expression and Its Association with GaM Sensitivity in 2D and 3D Culture Systems

Dose–response analyses revealed apparent differences in GaM cytotoxicity between 2D and 3D culture formats across the tested glioblastoma cell lines (Fig. 1). In 2D cultures, IC50 values ranged widely among cell lines. In contrast, in 3D cultures, a general shift toward higher drug tolerance was observed, with some lines showing pronounced resistance at clinically relevant concentrations. Specific lines (e.g., A-172, 3019, 3048) displayed a steeper dose–response curve in 3D compared with 2D conditions, indicating enhanced protection conferred by the 3D environment.

**Figure 1.**
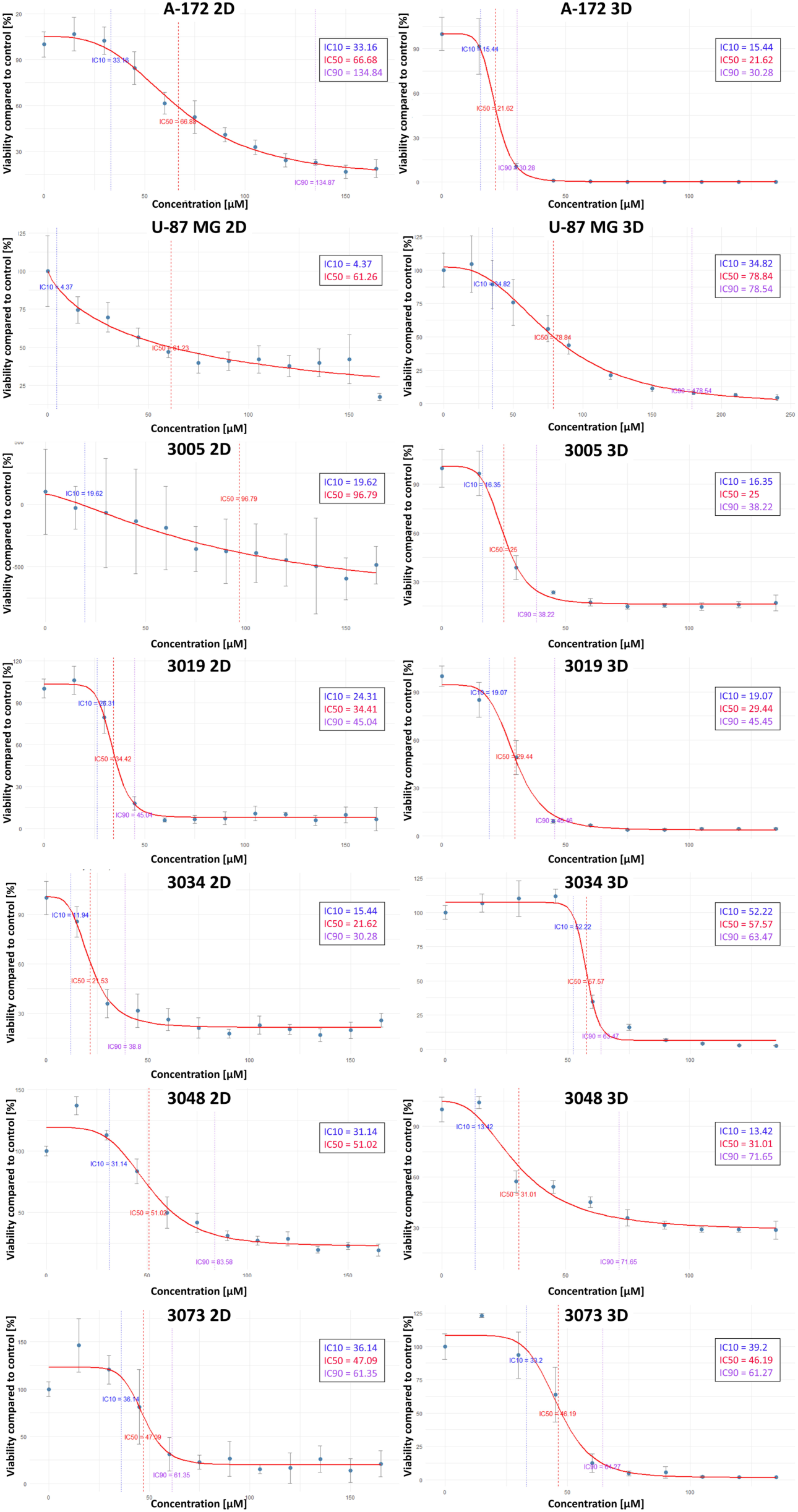
Percentage of cell viability compared to control after 72h incubation with GaM (15-165 µM – 2D, left; 15-135 µM -3D, right).

Basal TFRC expression levels differed substantially between 2D and 3D cultures (Fig. 2). Several cell lines (3005, 3019, 3034, 3073, U-87 MG) exhibited significantly higher TFRC protein abundance in 3D compared with 2D conditions, suggesting that culture dimensionality impacts iron metabolism–related pathways. Conversely, A-172 cells showed minimal differences between formats.

**Figure 2.**
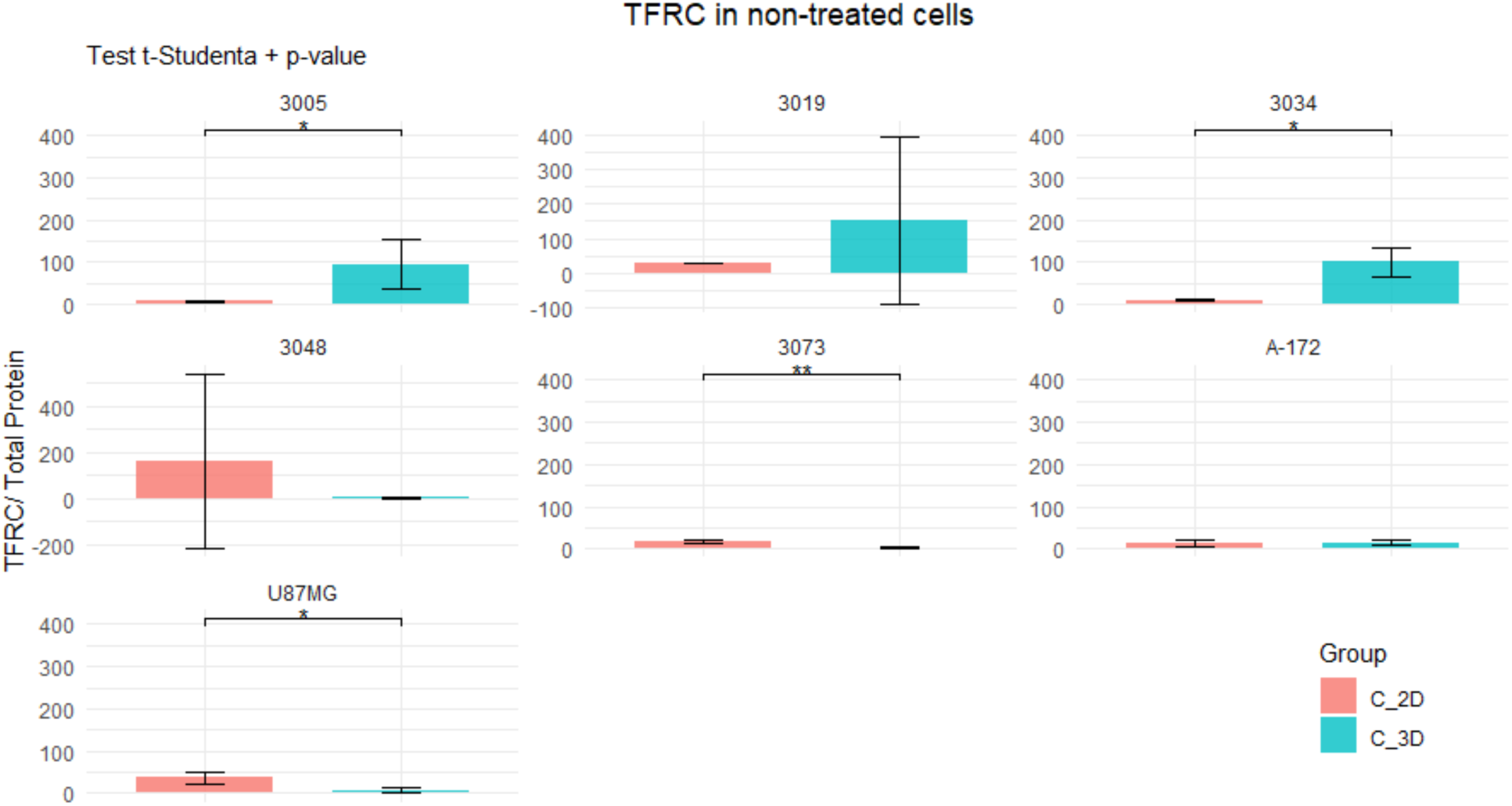
TFRC normalized for total protein content in non-treated control cells cultured in 2D and 3D format.

Correlation analysis further revealed that TFRC expression was positively associated with GaM sensitivity in 2D cultures, with a strong correlation observed between TFRC levels and IC50 values (R = 0.82, p = 0.024; Fig. 3). A similar but weaker, non-significant trend was seen for IC10 values (R = 0.39, p = 0.42). By contrast, no meaningful associations were detected in 3D cultures, where TFRC levels did not correlate with IC50 (R = 0.11, p = 0.82) or IC10 (R = –0.07, p = 0.87). However, the U-87 MG cell line behaved differently under 3D conditions. Excluding it as an outlier strengthened the Pearson correlation, yielding IC10 (R = 0.38, p = 4.45) and IC50 (R = 0.92, p = 0.01).

**Figure 3.**
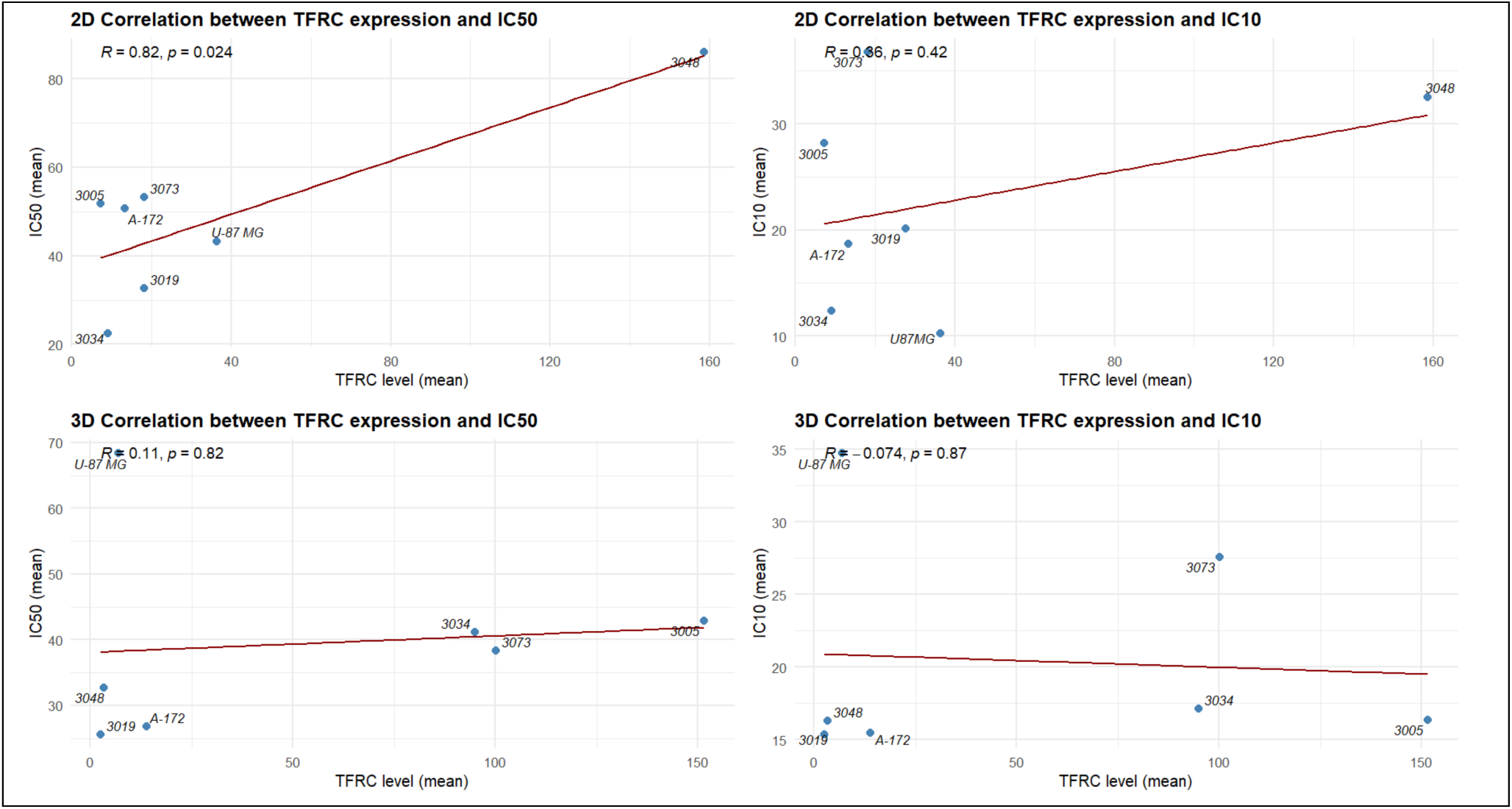
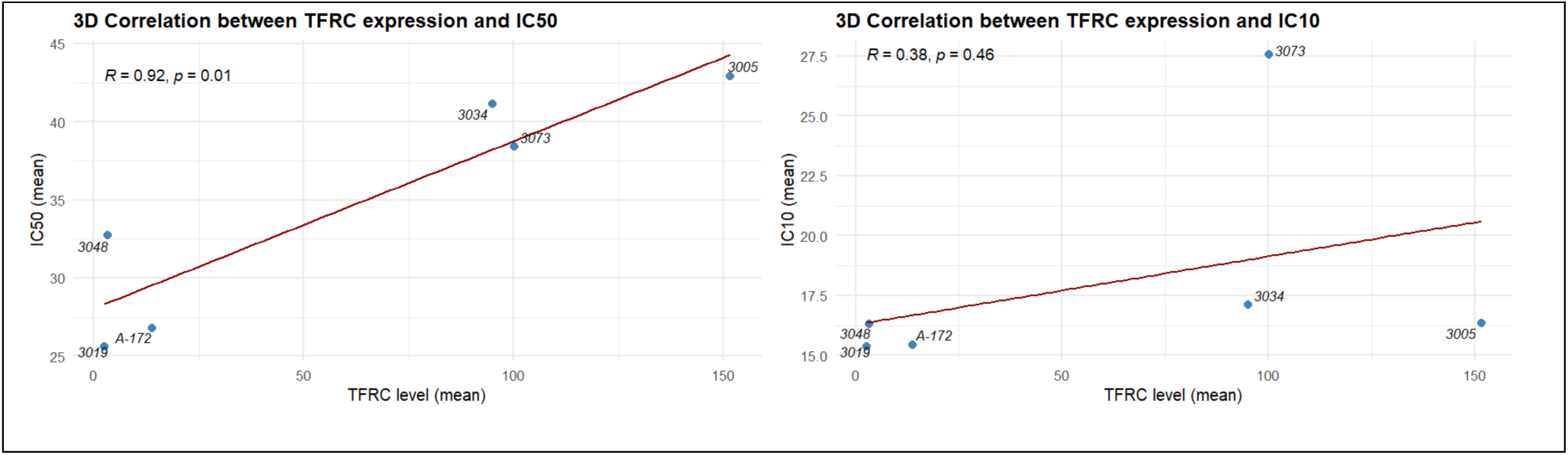
Correlation of detected TFRC level and IC50 and IC10 concentration determined with MTT (2D - top) and CellTiter Glo (3D - middle) assay and 3D correlation withoul U87MG (bottom).

### 3.2. Effects of GaM on TFRC Expression and Mitochondrial Respiration in 2D and 3D Culture Models

GaM treatment induced marked and cell line–dependent alterations in TFRC expression, with distinct responses observed between 2D and 3D culture formats (Fig. 1). In 2D conditions, a significant drop of TFRC level was observed in several lines (e.g., 3019, 3073, A-172) following 24h or 72h of exposure. In contrast, other lines (e.g., 3005, 3048) displayed variable or non-significant changes. By contrast, in 3D cultures, TFRC modulation was more heterogeneous: U-87 MG and 3034 exhibited a pronounced decrease after treatment, while 3073 demonstrated a transient upregulation at 72h, and 3019 showed no significant reduction even at 168h. These findings highlight both format-specific and temporal differences in the regulation of iron uptake pathways upon GaM exposure. Consistent with the TFRC data, OCR measurements revealed that GaM also differentially influenced mitochondrial respiration across culture formats (Fig. 2). In A-172, U-87 MG, 3073, and 3048 cells, GaM treatment led to a pronounced suppression of oxygen consumption, most evident under 3D conditions, suggesting impaired mitochondrial activity. In contrast, 3019 and 3005 cells maintained relatively stable OCR profiles irrespective of treatment, reflecting a higher metabolic resilience. Interestingly, 3034 cells displayed sustained OCR in 3D despite TFRC downregulation, indicating a possible shift to alternative metabolic pathways to support respiration. Taken together, these results demonstrate that GaM exerts a dual effect on glioblastoma cells by modulating TFRC expression and impairing mitochondrial respiration, with both responses being strongly dependent on cell line identity and the dimensionality of the culture system.

### 3.3. Multivariate Analysis of Metabolic Profiles in 2D and 3D Cultures

To investigate global metabolic alterations induced by GaM treatment, principal component analysis (PCA) (Fig. S2, Tab. S2) and partial least squares discriminant analysis (PLS-DA) were performed for each cell line under different experimental conditions (Fig. 6). Clear separation between sample groups was observed, with the degree of clustering varying according to cell line, treatment duration, and culture format.

**Figure 4.**
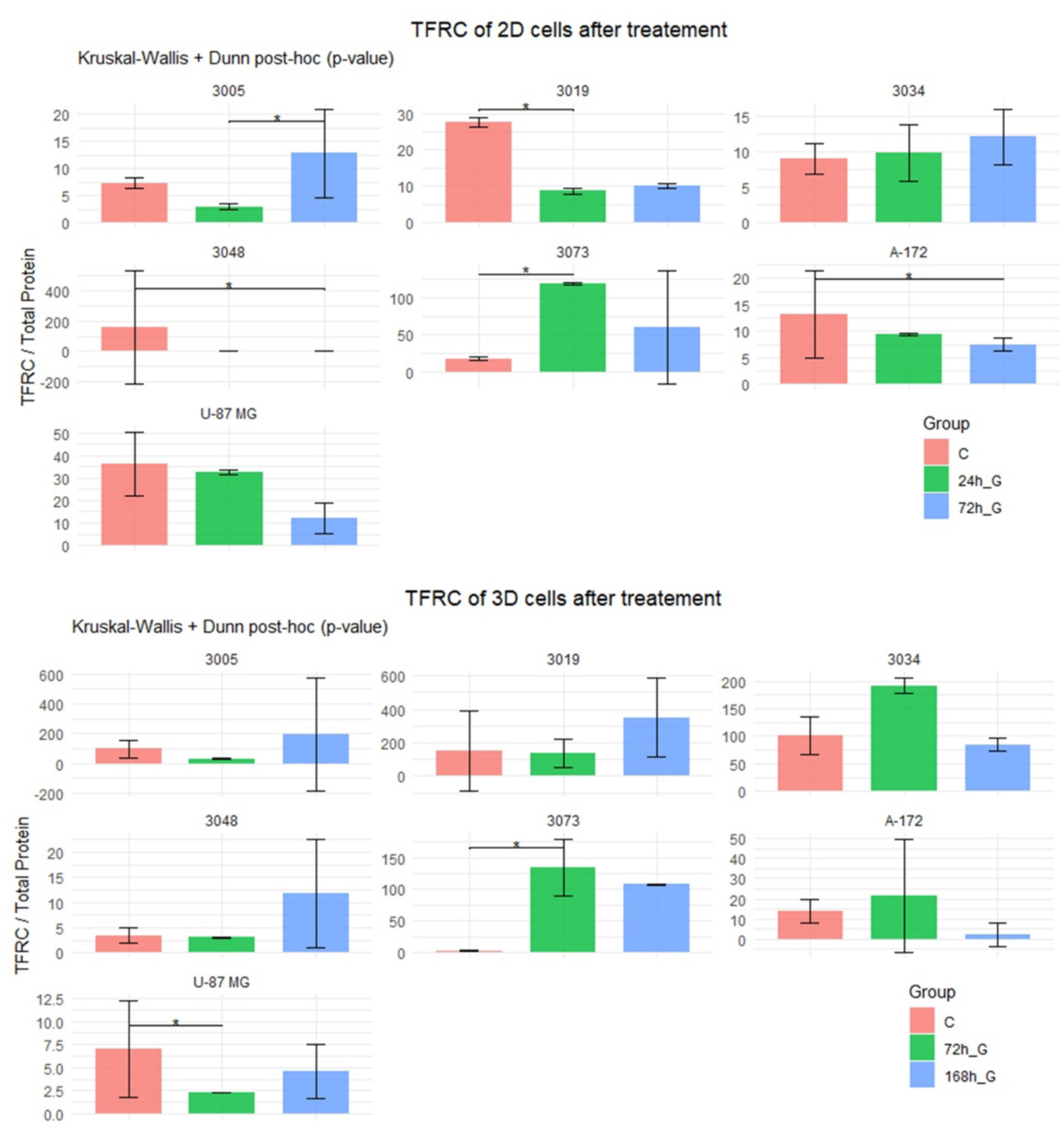
FTCR TFRC normalized total protein content in 2D cells (top) treated with IC50 concentration of GaM for 24h and 72h, and untreated control; 3D (bottom) treated with 2D IC50 concentration for 72h and 168h, and untreated control.

**Figure 5.**
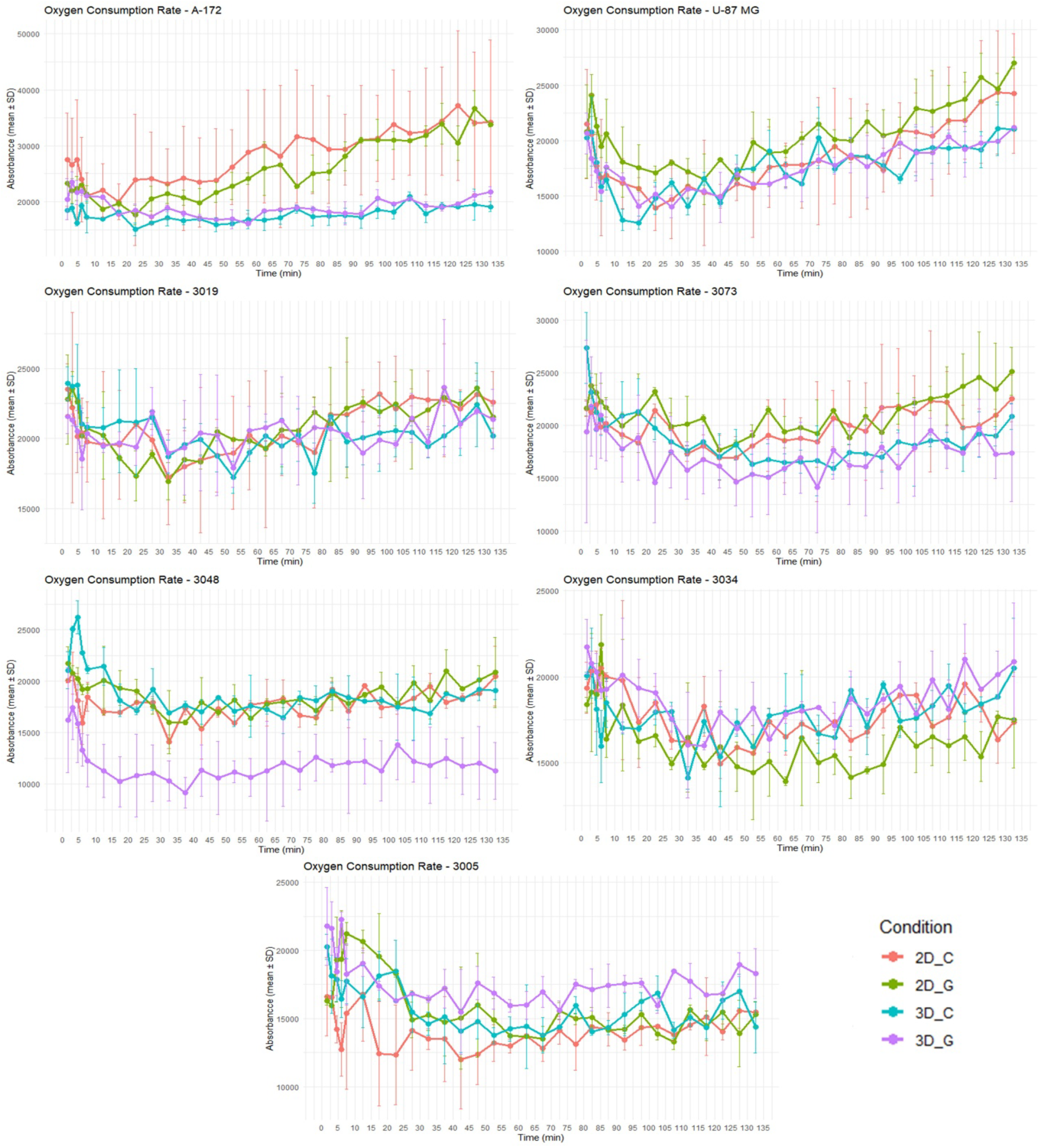
OCR level in 2D and 3D culture immediately after 2DIC50 concentration and untreated control.

**Figure 6.**
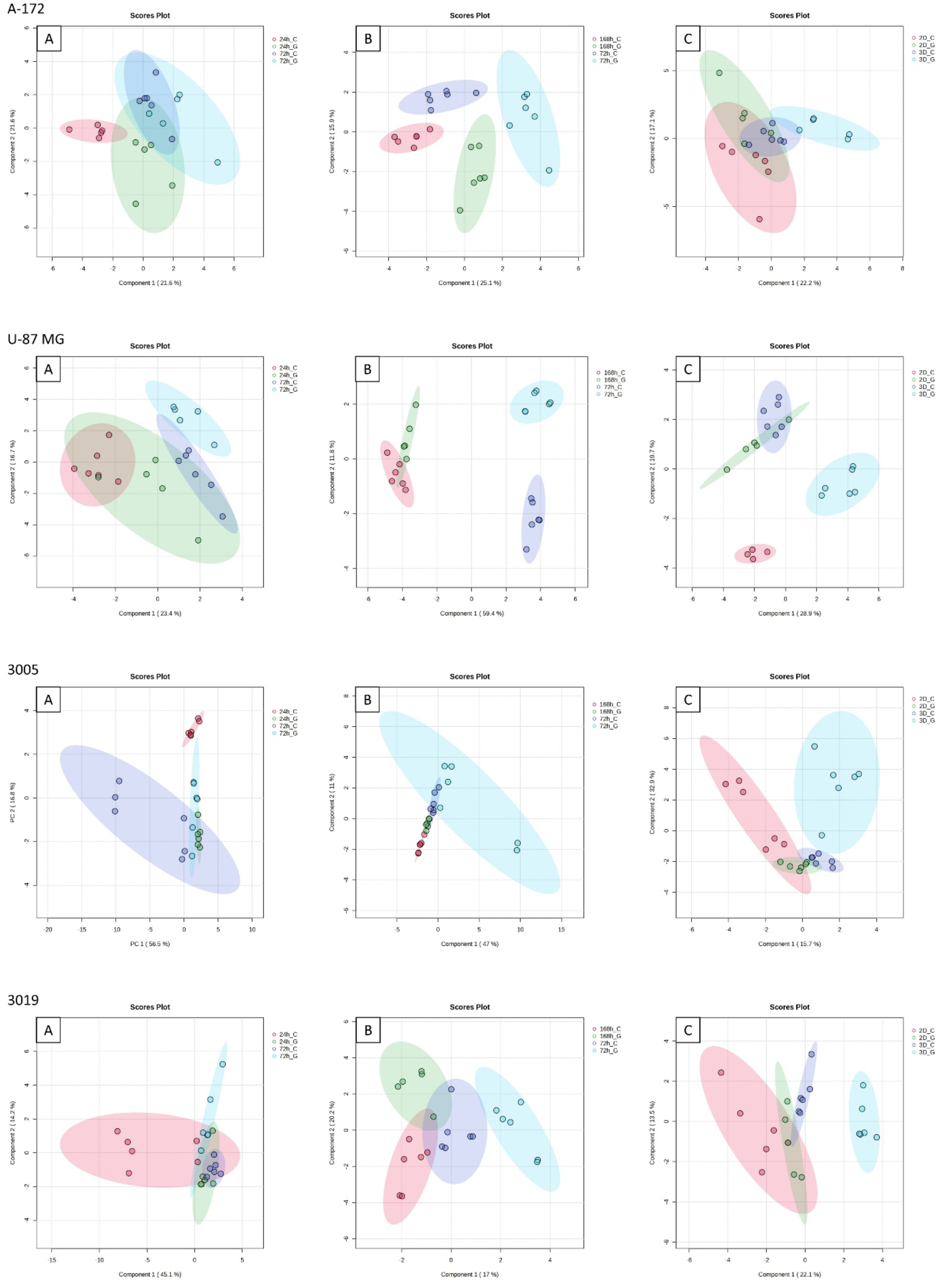
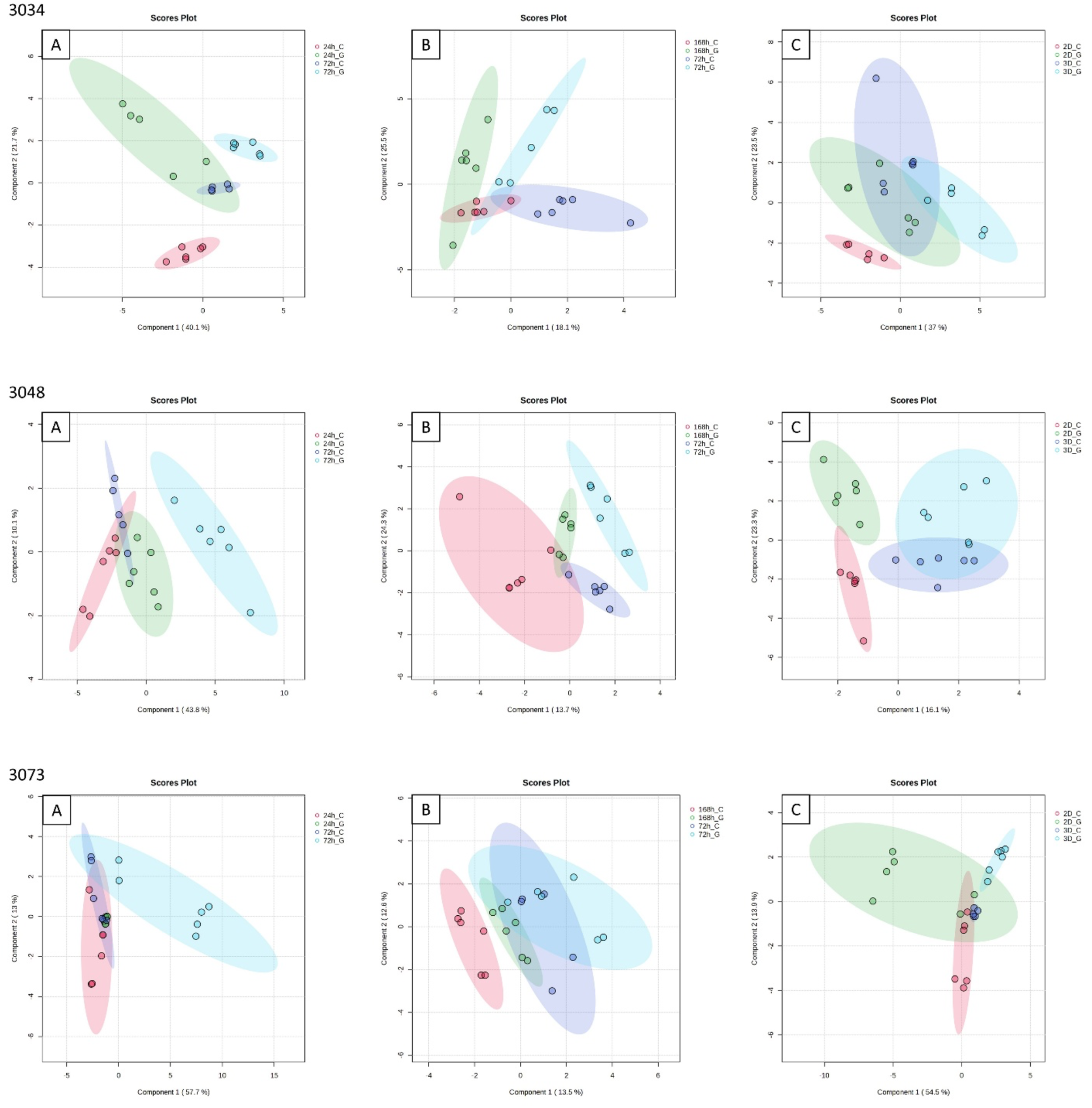
PLS-DA score plots showing separation of all cell lines in A) 2D 24h vs 72h, B) 3D 72h vs 168h, C) 72h 2D vs 3D (n=6).

In A-172 and U-87 MG cells, temporal effects were the predominant drivers of variance, with early (24 h) and late (72–168 h) treatment groups forming distinct clusters. Treatment-dependent separation was also evident, particularly at later time points, although overlap between control and GaM-exposed samples persisted in some comparisons. Among the patient-derived glioblastoma models, 3005 and 3048 exhibited a stronger treatment-dependent response, as GaM exposure led to pronounced segregation of metabolic profiles from untreated controls, especially under 3D conditions. In contrast, 3019 and 3034 cells demonstrated striking format-specific clustering, with 2D and 3D cultures forming distinct groups regardless of treatment, underscoring the dominant effect of dimensionality in shaping metabolic programs. Finally, 3073 cells displayed combined influences of both time and culture format, with 3D groups clearly separated from their 2D counterparts and temporal clustering evident within each format.

These analyses indicate that GaM induces robust and cell line–specific metabolic reprogramming. In some lines (e.g., 3005, 3048), treatment was the dominant factor, whereas in others (e.g., 3019, 3034) the culture format exerted the most decisive influence. In contrast, for A-172, U-87 MG, and 3073, metabolic variation was driven primarily by treatment duration.

The comparative analysis of treated cells and untreated control within one incubation time further explored the metabolomic separation upon GaM exposure (Fig S3-S9, Tab S1). Targeted metabolomic profiling revealed consistent alterations in several metabolites across glioblastoma models following GaM exposure (Tab. S3). Among the significantly perturbed metabolites, uracil accumulated in multiple models, indicating disruption of pyrimidine metabolism, which may reflect altered nucleotide turnover or stress-related RNA degradation (Fig.7 and 8). In turn, tryptophan levels were markedly reduced in treated cells, suggesting interference with tryptophan catabolism and potentially implicating the kynurenine pathway (Fig. 9 and 10). Methionine levels were significantly diminished upon treatment, consistent with impaired one-carbon metabolism and reduced methylation potential (Fig. S10 and S11), while allantoin, a marker of oxidative stress and purine catabolism, was strongly elevated, reflecting treatment-induced redox imbalance (Fig. S12 and S13). All scope of significantly changed metabolites can be found in Table S2.

**Figure 7.**
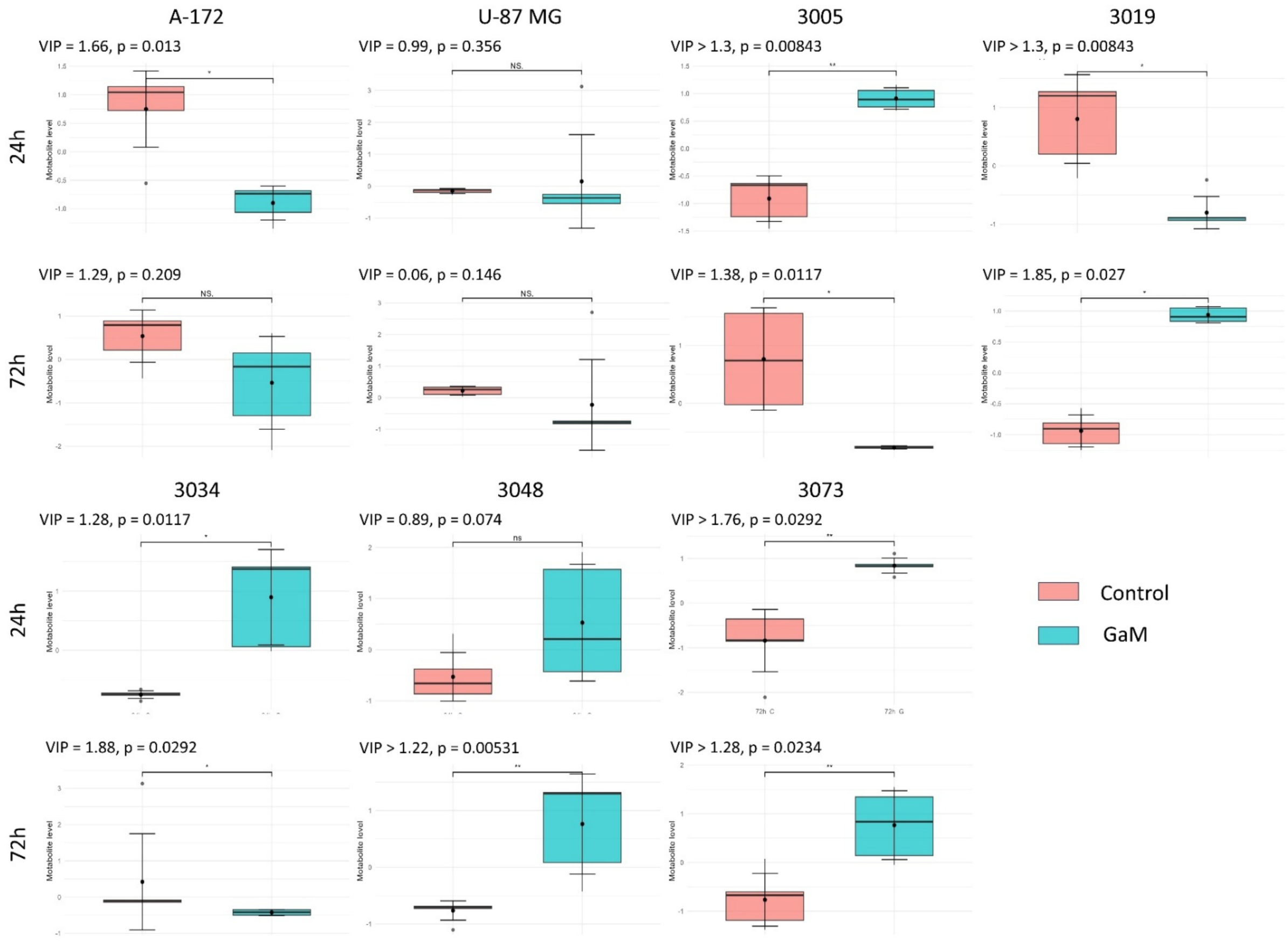
Change in levels of uracil between GaM treated cells and untreated control in 2D culture with VIP score and p-value (FDR).

**Figure 8.**
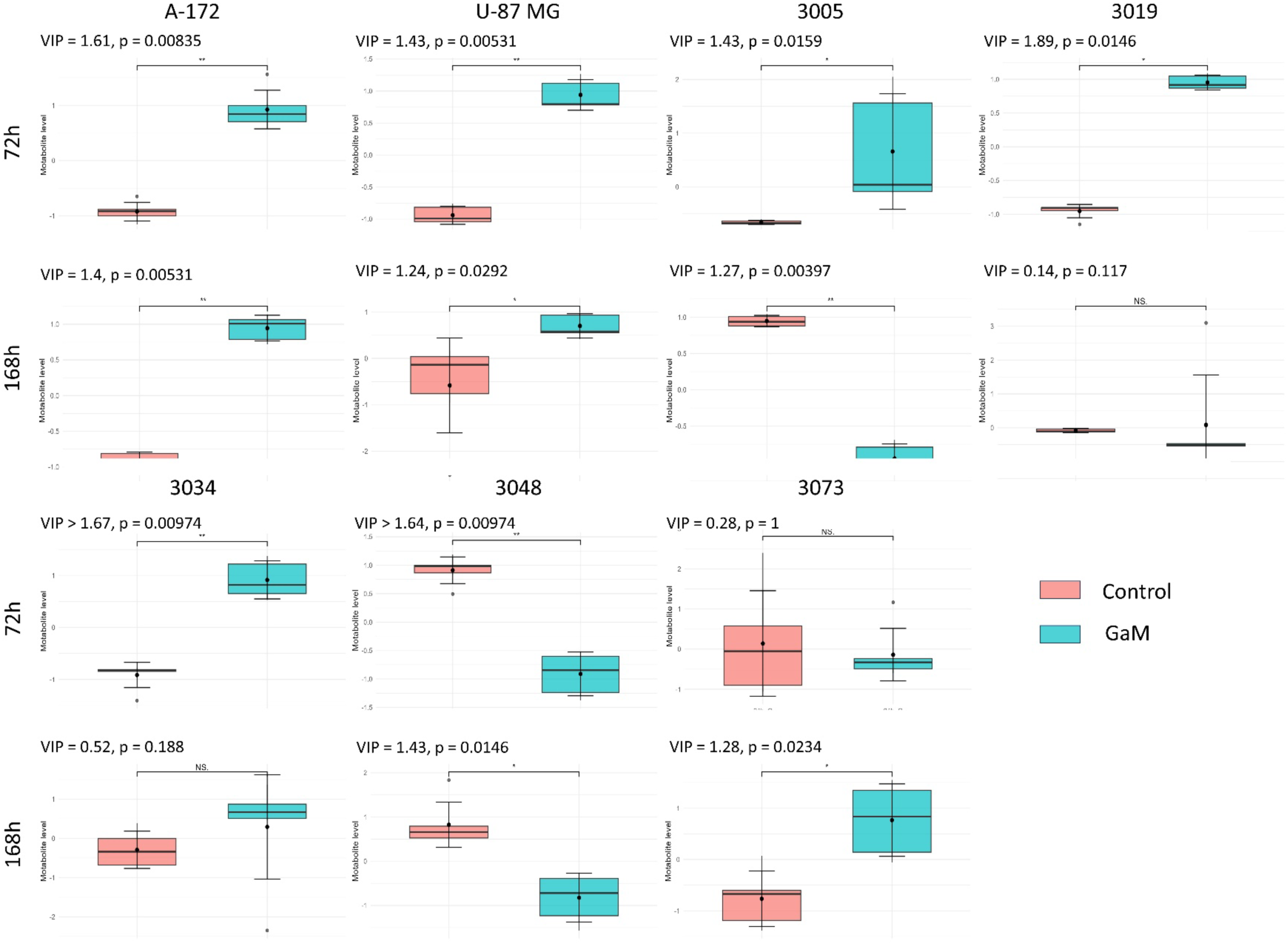
Change in levels of uracil between GaM treated cells and untreated control in 3D culture with VIP score and p-value (FDR).

**Figure 9.**
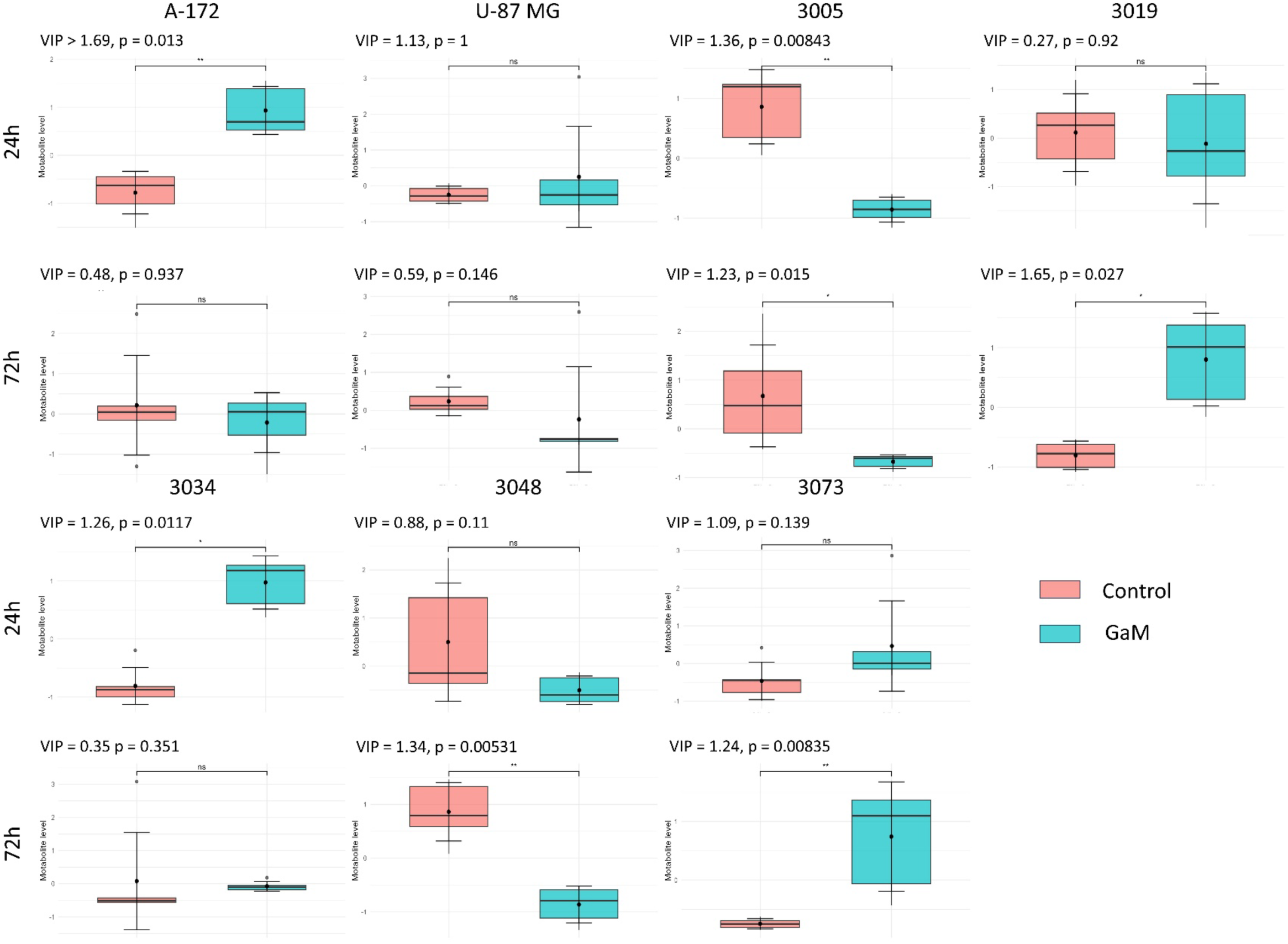
Change in levels of tryptophan between GaM treated cells and untreated control in 2D culture with VIP score and p-value (FDR).

**Figure 10.**
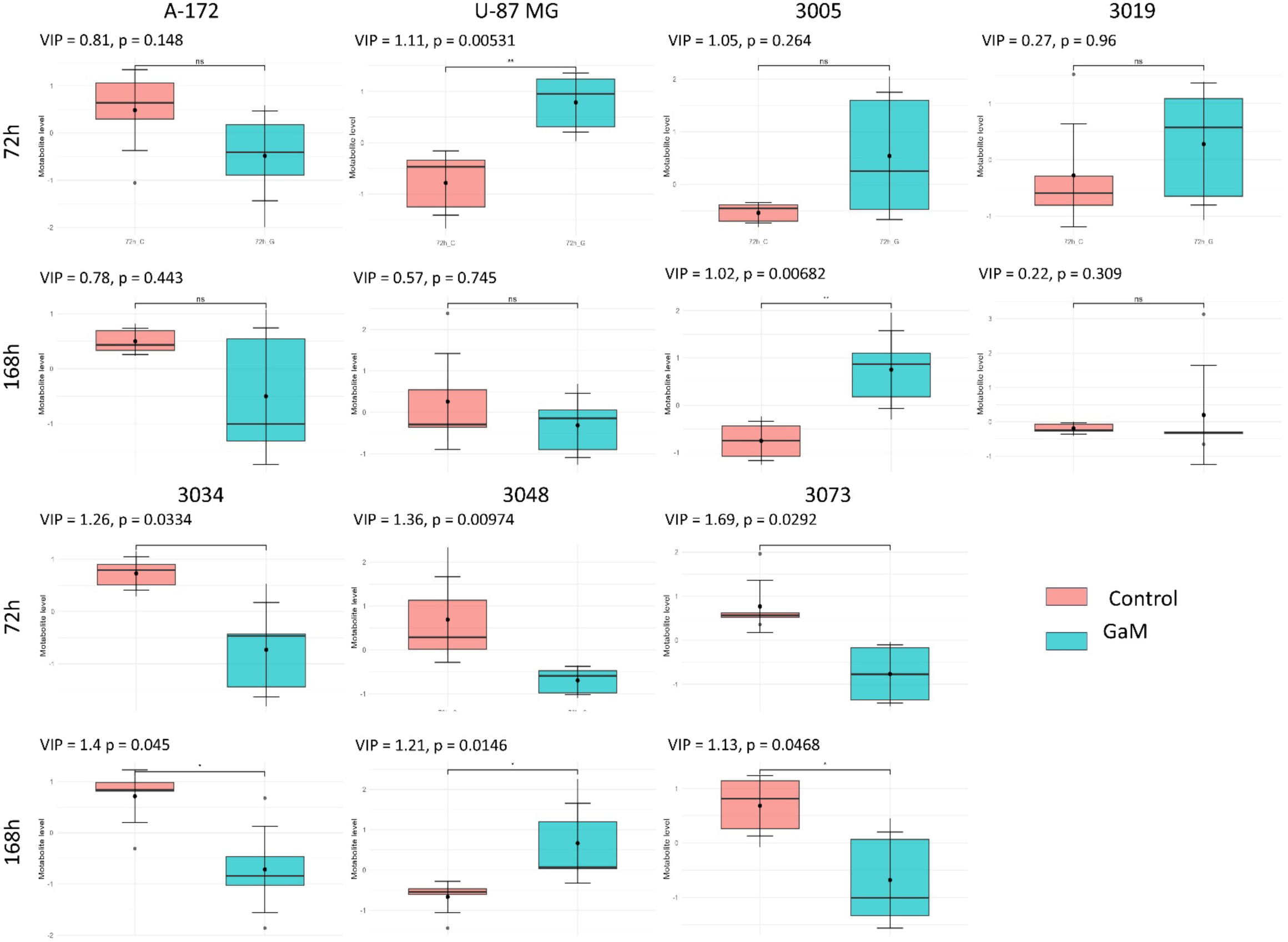
Change in levels of tryptophan between GaM treated cells and untreated control in 2D culture with VIP score and p-value (FDR).

Pathway enrichment analysis supported these observations, highlighting significant impacts on amino acid metabolism (tryptophan, methionine), nucleotide metabolism (uracil, purines), and redox-related pathways (allantoin) (Fig. 11, Table 1). These pathways emerged among the most significantly perturbed, with high pathway impact scores, suggesting that GaM treatment broadly reprograms metabolic networks essential for glioblastoma proliferation and survival.

**Figure 11.**
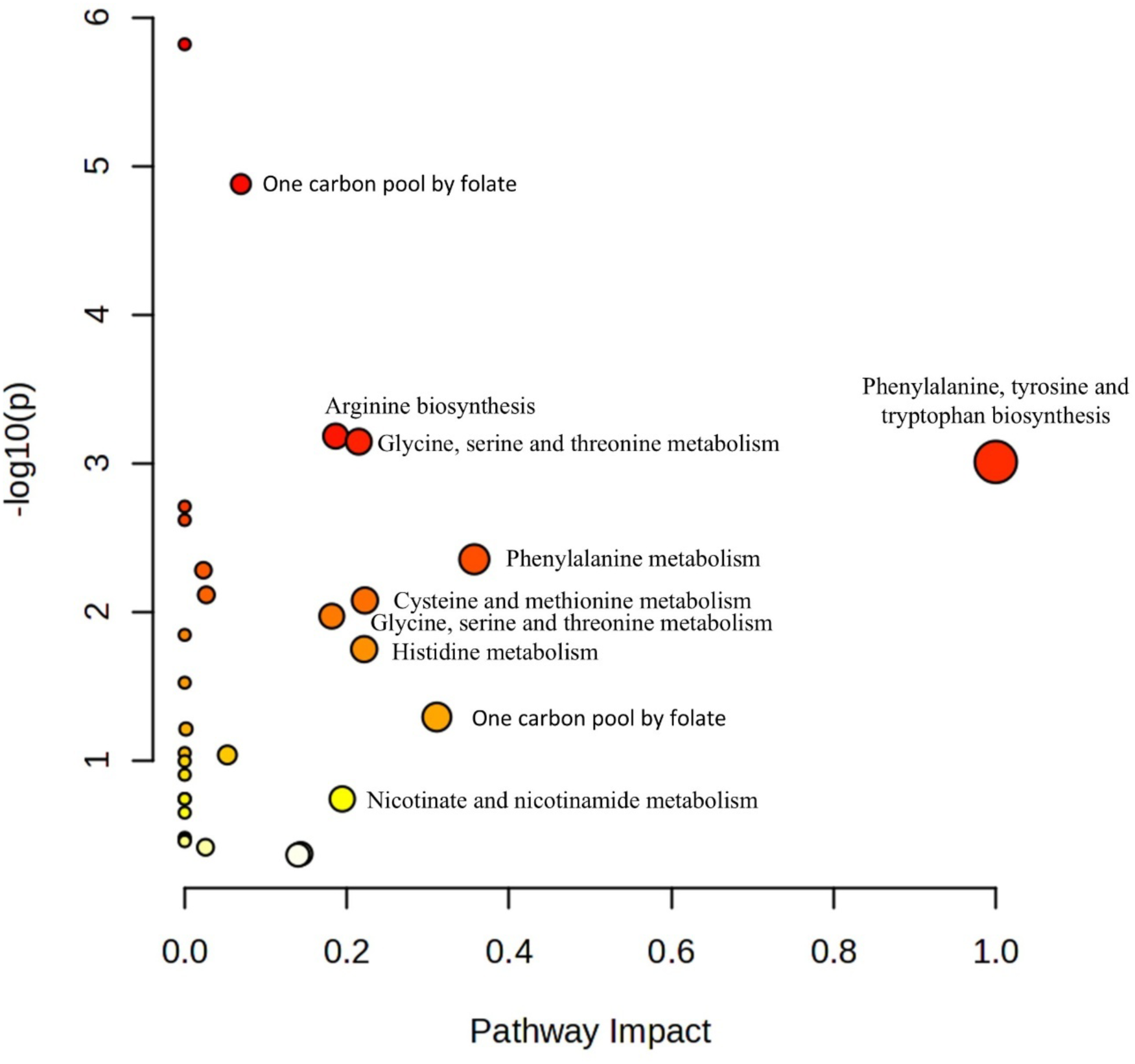
Pathway analysis of significantly differential features from metabolomic analysis.

**Table 1.** Detailed information on Pathway Analysis performed based on metabolites of VIP score > 1 and FDR < 0.05.

| Pathway Name | Match Status | p | -log(p) | Holm p | FDR | Impact |
| --- | --- | --- | --- | --- | --- | --- |
| Valine, leucine, and isoleucine biosynthesis | 4/8 | 1.23E-06 | 5.9096 | 9.85E-05 | 9.85E-05 | 0 |
| One carbon pool by folate | 5/26 | 1.32E-05 | 4.8783 | 0.001045 | 0.000529 | 0.06931 |
| Arginine biosynthesis | 3/14 | 0.000566 | 3.2473 | 0.044133 | 0.015088 | 0.18617 |
| Glycine, serine, and threonine metabolism | 4/33 | 0.000715 | 3.1455 | 0.055082 | 0.014307 | 0.21459 |
| Phenylalanine, tyrosine, and tryptophan biosynthesis | 2/4 | 0.000887 | 3.0523 | 0.068271 | 0.017733 | 1 |
| Pantothenate and CoA biosynthesis | 3/20 | 0.001689 | 2.7725 | 0.12832 | 0.027015 | 0 |
| Nitrogen metabolism | 2/6 | 0.002183 | 2.6609 | 0.16375 | 0.02911 | 0 |
| Phenylalanine metabolism | 2/8 | 0.004014 | 2.3964 | 0.29706 | 0.045496 | 0.35714 |
| Glutathione metabolism | 3/28 | 0.00455 | 2.342 | 0.33212 | 0.045496 | 0.02309 |
| Glyoxylate and dicarboxylate metabolism | 3/32 | 0.00667 | 2.1759 | 0.48025 | 0.052934 | 0.02667 |
| Cysteine and methionine metabolism | 3/33 | 0.007278 | 2.138 | 0.51677 | 0.052934 | 0.22222 |
| Arginine and proline metabolism | 3/36 | 0.009298 | 2.0316 | 0.64156 | 0.061986 | 0.18139 |
| Valine, leucine, and isoleucine degradation | 3/40 | 0.012459 | 1.9045 | 0.84721 | 0.076671 | 0 |
| Histidine metabolism | 2/16 | 0.016197 | 1.7906 | 1 | 0.092555 | 0.22131 |
| beta-Alanine metabolism | 2/21 | 0.0273 | 1.5638 | 1 | 0.1456 | 0 |
| Alanine, aspartate, and glutamate metabolism | 2/28 | 0.04663 | 1.3313 | 1 | 0.23315 | 0.3109 |
| Purine metabolism | 3/70 | 0.054267 | 1.2655 | 1 | 0.25538 | 0.00158 |
| Pyrimidine metabolism | 2/39 | 0.084215 | 1.0746 | 1 | 0.35725 | 0.05261 |
| Thiamine metabolism | 1/7 | 0.084848 | 1.0714 | 1 | 0.35725 | 0 |
| Taurine and hypotaurine metabolism | 1/8 | 0.096396 | 1.0159 | 1 | 0.38558 | 0 |
| Biotin metabolism | 1/10 | 0.11908 | 0.92417 | 1 | 0.45363 | 0 |
| D-Amino acid metabolism | 1/15 | 0.17344 | 0.76086 | 1 | 0.57813 | 0 |
| Butanoate metabolism | 1/15 | 0.17344 | 0.76086 | 1 | 0.57813 | 0 |
| Nicotinate and nicotinamide metabolism | 1/15 | 0.17344 | 0.76086 | 1 | 0.57813 | 0.1943 |
| Ubiquinone and other terpenoid-quinone biosynthesis | 1/19 | 0.21462 | 0.66834 | 1 | 0.68677 | 0 |
| Lysine degradation | 1/30 | 0.31805 | 0.49751 | 1 | 0.95829 | 0 |
| Porphyrin metabolism | 1/31 | 0.32678 | 0.48575 | 1 | 0.95829 | 0 |
| Sphingolipid metabolism | 1/32 | 0.3354 | 0.47443 | 1 | 0.95829 | 0 |
| Tryptophan metabolism | 1/41 | 0.40845 | 0.38886 | 1 | 1 | 0.14305 |
| Tyrosine metabolism | 1/42 | 0.41608 | 0.38083 | 1 | 1 | 0.13972 |

## 4. Discussion

GBM is an aggressive and incurable tumor of the central nervous system. The alarming increase in incidence, frequent recurrences, and high mortality has driven a steady rise in research on new and effective therapies in recent years. A next-generation compound, GaM, showed greater efficacy in preclinical studies compared with GaN against hepatocellular carcinoma cells and GaN-resistant lymphoma cells, suggesting that the mechanism of gallium ion transport in complex with maltolate differs from that of GaN [43,44]. The proven effectiveness of GaM against treatment-resistant cells provides strong prospects for developing an effective therapy for aggressive and incurable GBM.

The present study aimed to evaluate the cytotoxic activity of GaM against GBM cells in both 2D and 3D culture models. Our results indicate that GaM perturbs iron-dependent metabolism in GBM, coherently linking TFRC biology, mitochondrial respiration (OCR), and multivariate shifts in metabolite profiles. GaM exploits transferrin, a protein responsible for iron transfer into the cell, to access the brain and GBM cells. Chitambar et al. described that GaM disrupts mitochondrial function (notably complex I via impaired iron–sulfur cluster assembly) and inhibits the iron-dependent RRM2 subunit of ribonucleotide reductase—effects demonstrated in U-87 MG/D54 cells and validated *in vivo* where GaM retards GBM growth and alters iron markers^12^. Glioblastoma cell lines have been shown to overexpress transferrin receptors frequently, not only TfR1; TfR2 is highly and commonly expressed in GBM, it has been associated with grade and proliferation, underscoring that the phenotype of iron transfer into the cells is broader and may shape sensitivity to iron-mimetic therapies^27^.

Culture dimension emerged as a dominant modifier of GaM response in our models. Several lines (A-172, U-87 MG, 3073, 3048) showed stronger OCR suppression and more apparent metabolic separation in 3D than 2D, consistent with evidence that advanced 3D GBM systems better reproduce diffusion barriers, ECM/mechanical cues, chemical gradients, and BBB contributions—features that modulate drug penetration and frequently reduce apparent drug sensitivity relative to monolayers^28^. In particular, an engineered human BBB–GBM co-culture showed decreased temozolomide sensitivity with increased tumor spatial organization and BBB involvement, exemplifying how microenvironmental architecture can decouple single-target predictors (e.g., TFRC levels) from whole-cell pharmacologic outcomes.

Model identity also influenced GaM phenotypes. Established lines (A-172, U-87 MG) tended to display time-dependent separation and robust OCR effects, whereas multiple patient-derived lines showed either format-dominated clustering (2D vs 3D outweighing treatment; e.g., 3019, 3034) or pronounced treatment-specific separation (e.g., 3005, 3048). This mirrors comparisons between established GBM lines and patient-derived/neurosphere models, where 3D states rewire metabolism and drug sensitivity^29^. These considerations support prioritizing 3D and patient-derived platforms for GaM evaluation and for combination-strategy testing.

The observed alterations in tryptophan, uracil, methionine, and allantoin provide insights into the cellular response to GaM treatment. Tryptophan depletion is particularly relevant, as it may reflect increased catabolism through the kynurenine pathway, a route tightly linked to immunosuppression and tumor progression ^30,31^. Reduced availability of tryptophan could therefore impair protein synthesis and alter immune interactions, suggesting that GaM interferes with both metabolic and signalling roles of this essential amino acid—similarly, the consistent changes in uracil point to disrupted pyrimidine metabolism. Accumulation of uracil has been associated with imbalances in nucleotide pools and misincorporation into DNA, which requires base excision repair. Such changes may contribute to replication stress and reduced proliferation under GaM exposure^32,33^. The decrease in methionine levels further underscores the influence on DNA. Methionine is a key donor in one-carbon metabolism and methylation reactions, and its depletion suggests impaired DNA and histone methylation capacity, which could translate into epigenetic instability and altered gene regulation in glioblastoma cells ^34,35^.

Finally, the significant increase in allantoin reflects perturbations in purine metabolism and elevated oxidative stress. Allantoin accumulation is a recognized marker of reactive oxygen species (ROS) activity. This implies that GaM treatment may induce redox imbalance and oxidative damage, compromising glioblastoma cell viability^36,37^. These findings suggest that GaM exerts its effects through a multifaceted disruption of amino acid, nucleotide, and redox metabolism. This combination of metabolic stressors likely contributes to impaired biosynthesis, genomic instability, and oxidative damage, thereby sensitizing glioblastoma cells to treatment.

Summarizing, at the metabolite level, methionine, uracil, and allantoin provide a concise, mechanism-anchored narrative that aligns with our pathway analysis. Tryptophan depletion implicates kynurenine/immune-metabolic axes; methionine decrease suggests pressure on one-carbon metabolism and methylation capacity; uracil dysregulation points to pyrimidine pool imbalance and nucleic acid turnover stress; and allantoin elevation is compatible with ROS-linked purine oxidation—all converging with our OCR data on mitochondrial compromise under GaM^12^. Together, these data support amino-acid, nucleotide, and redox stress as integrated drivers of GaM cytotoxicity.

Translationally, GaM has shown in vivo activity in GBM xenografts, including oral delivery that slows tumor growth and extends disease-specific survival^38^; early reports also noted reductions in tumor and relative cerebral blood volume, and continuous administration suppressed glioma growth and *in vivo*^12^. These findings along with literature on gallium complexes as anticancer agents reinforce the rationale for GaM development and for integrating metabolic biomarkers (e.g., TFRC and our four-metabolite panel) into preclinical–clinical translation^8^. Finally, synergy with complex-I inhibition (e.g., metformin) in 2D and 3D GBM further supports a mitochondria-centric vulnerability under GaM treatment that may be exploitable in rational combinations^39^.

Together, these data argue that 3D, patient-derived systems provide a more predictive test bed for GaM than conventional monolayers and that integrated OCR/metabolomics readouts can serve as practical pharmacodynamic markers. However, a study with a comprehensive patient-derived glioblastoma cell panel should be performed, further exploring the role of the *in vitro* microenvironment. Direct quantification of gallium uptake and GaM penetration into the 3D spheroid could provide more information on the real toxicity of the drug, improving *in vitro-in vivo* extrapolation of results. Our findings support further development of GaM for GBM and provide a framework for selecting biomarkers and model systems that better forecast clinical performance. Conclusion

Culture dimensionality significantly determines gallium maltolate (GaM) response in glioblastoma. Across established (A-172, U-87 MG) and patient-derived lines (3005, 3019, 3034, 3048, 3073), GaM reduced viability and suppressed mitochondrial respiration, with a consistent rightward shift of dose–response and more potent OCR inhibition in 3D than in 2D. TFRC levels associated with GaM sensitivity in 2D but not 3D indicated context-dependent biomarker performance. Metabolomics and multivariate analyses converged on a compact signature— tryptophan, methionine, uracil, and allantoin—consistent with stress in amino-acid, one-carbon/nucleotide, and redox pathways alongside mitochondrial dysfunction. These findings support using 3D, patient-derived systems with integrated OCR/ metabolomics as more predictive platforms for GaM evaluation and rational combination strategies.

## 5. Associated content

Supplementary material includes:

Figure S1. Principal component analysis (PCA) score plots of all analyzed samples and extraction quality control (QC) samples.

Figure S2. PCA score plots showing separation of all cell lines in A) 2D 24h vs 72h, B) 3D 72h vs 168h, C) 72h 2D vs 3D (n=6).

Figutre S3. PCA (top) and PLS-DA scrore plots showing separations of A-172 cell line in 2D and 3D, treated (G) and untreated (C).

Figutre S4. PCA (top) and PLS-DA scrore plots showing separations of U-87 MG cell line in 2D and 3D, treated (G) and untreated (C).

Figutre S5. PCA (top) and PLS-DA scrore plots showing separations of 3005 MG cell line in 2D and 3D, treated (G) and untreated (C).

Figutre S6. PCA (top) and PLS-DA scrore plots showing separations of 3019 MG cell line in 2D and 3D, treated (G) and untreated (C).

Figutre S7. PCA (top) and PLS-DA scrore plots showing separations of 3034 MG cell line in 2D and 3D, treated (G) and untreated (C).

Figutre S8. PCA (top) and PLS-DA scrore plots showing separations of 3048 MG cell line in 2D and 3D, treated (G) and untreated (C).

Figutre S9. PCA (top) and PLS-DA scrore plots showing separations of 3073 MG cell line in 2D and 3D, treated (G) and untreated (C).

Figure S10. Change in levels of methionine between GaM treated cells and untreated control in 2D culture with VIP score and p-value (FDR).

Figure S11. Change in levels of methionine between GaM treated cells and untreated control in 3D culture with VIP score and p-value (FDR).

Figure S12. Change in levels of allatoin between GaM treated cells and untreated control in 2D culture with VIP score and p-value (FDR).

Figure S13. Change in levels of allatoin between GaM treated cells and untreated control in 3D culture with VIP score and p-value (FDR).

Code S1. Dose-response curve with IC90, IC50 and IC10 calculations.

Code S2. T-test for TFRC level determination and significance of changes determination in 2D and 3D control.

Code S3. Pearson correlation test between IC10 and TFRC level in 2D and 3D cell lines.

Code S.4 TFRC level determination in treated cells and untreated control, Kurskal-Wallis test with dunn post-hoc.

Code S5. The Oxygen Consuption Rate curve in time.

Code S6 Wilcox test with FDR correction and selection of metabolites with VIP > 1 and adj. p-value (FDR).

Table S1. Cross-validated PLS-DA performance across GBM models.

Table S2. Comprehensive metabolite panel by cell line (3005, 3019, 3034, 3048, 3073, A-172, U-87 MG): VIP, FDR, stars

## 6. Acknowledgements

We thank Kamil Łuczykowski and Joanna Bogusiewicz for technical support with LC-MS workflows. We thank Natalia Warmuzińska for comprehensive statistical advice.

Portions of language editing were assisted by ChatGPT (OpenAI, GPT-5 Thinking, 16.09.2025).

## 7. Author contributions

The manuscript was written through the contributions of all authors. All authors have given approval to the final version of the manuscript

---

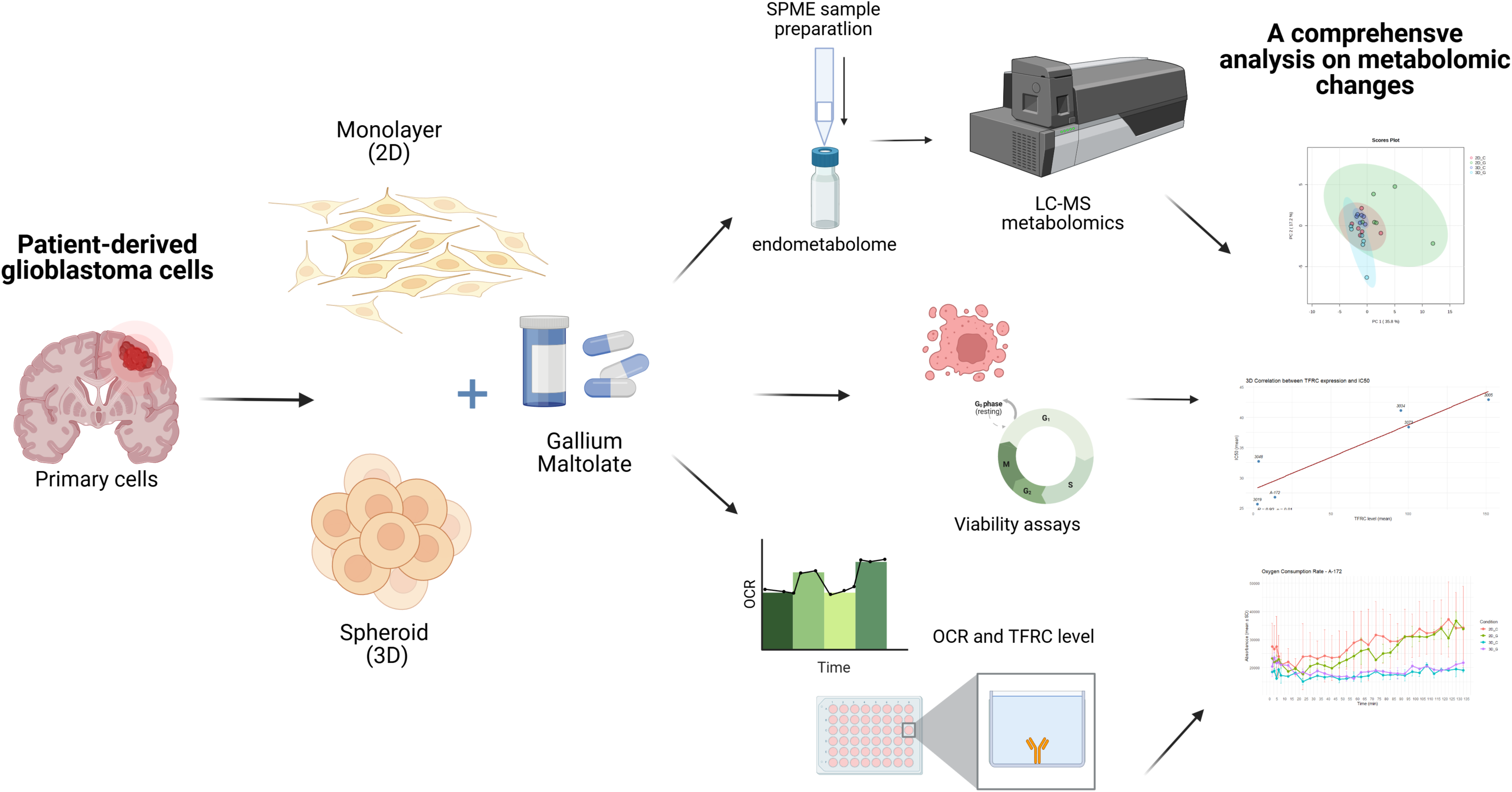

