## Supplementary materials for "Culture Dimensionality Governs Gallium Maltolate Response in Glioblastoma: Comparative Analyses in 2D and 3D Models"

Table S1. Basic characteristics of HGCC glioblastoma cells.

| Cell Line | HGCC_Subtype | Sex | Age (year) | Overall Survival (days) | Vital Status | Diagnosis | WHO Grade |
|---|---|---|---|---|---|---|---|
| U3005MG | Classical (CL) | Male | 65 | 26 | DEAD | Glioblastoma | IV |
| U3019MG | Proneural (PN) | Female | 82 | 117 | DEAD | Glioblastoma | IV |
| U3034MG | Mesenchymal (MS) | Male | 73 | 539 | DEAD | Glioblastoma | IV |
| U3048MG | Classical (CL) | Male | 77 | 279 | DEAD | Glioblastoma | IV |
| U3073MG | Mesenchymal (MS) | Male | 71 | 481 | DEAD | Glioblastoma | IV |

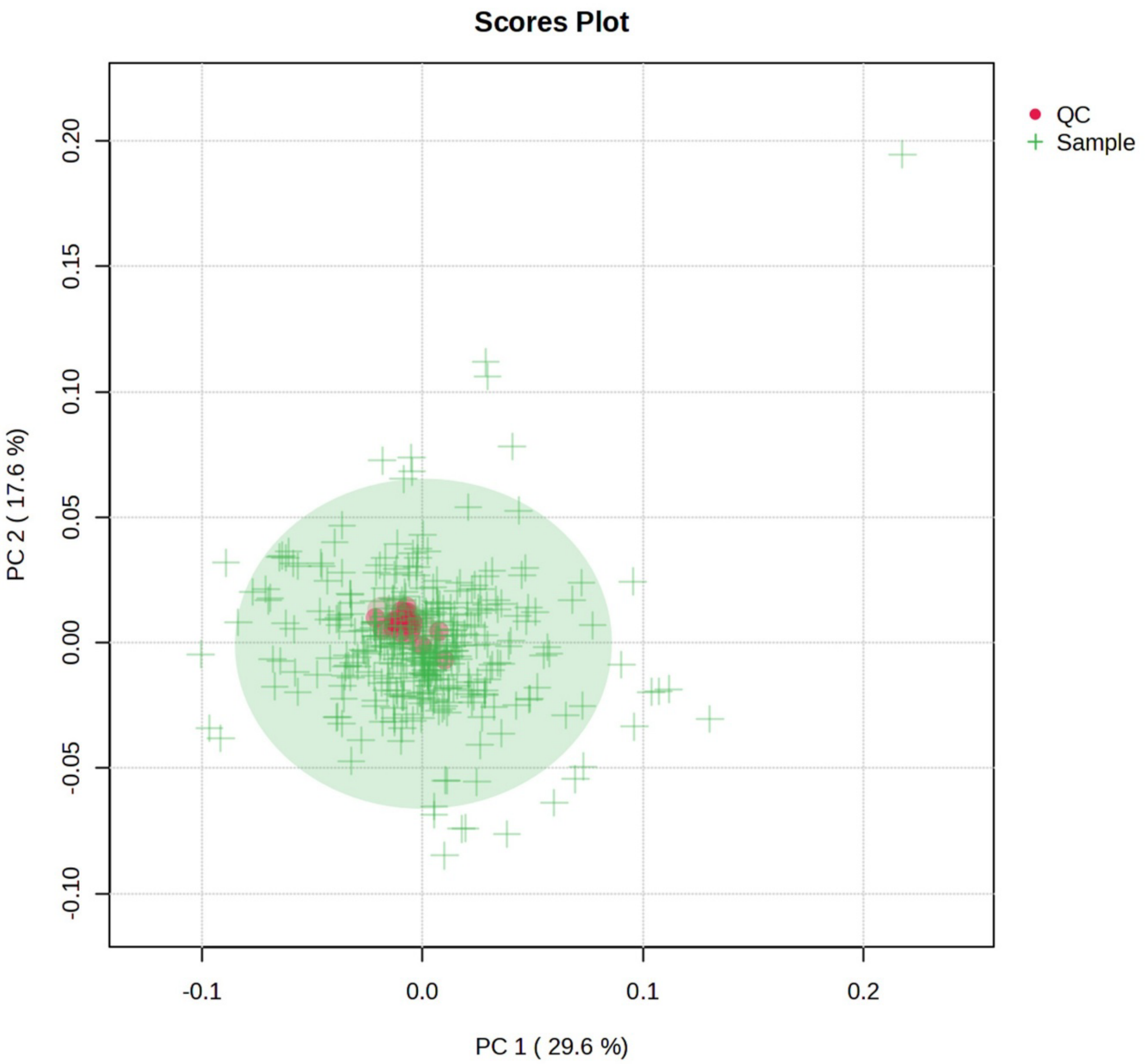

Figure S1. Principal component analysis (PCA) score plots of all analyzed samples and extraction quality control (QC) samples.

A-172

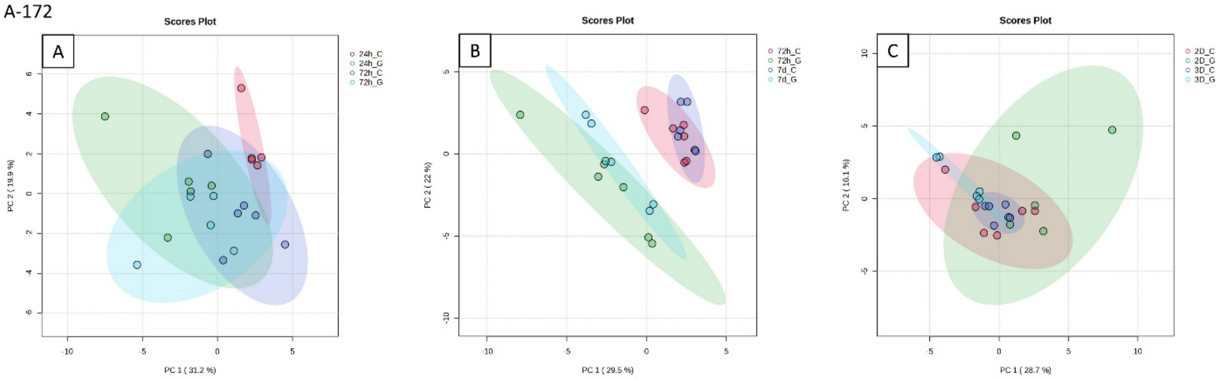

U-87 MG

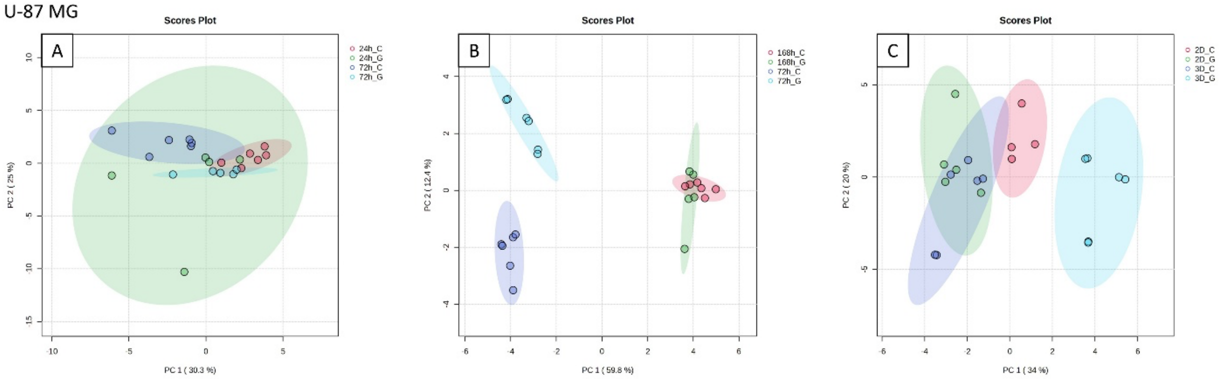

3005

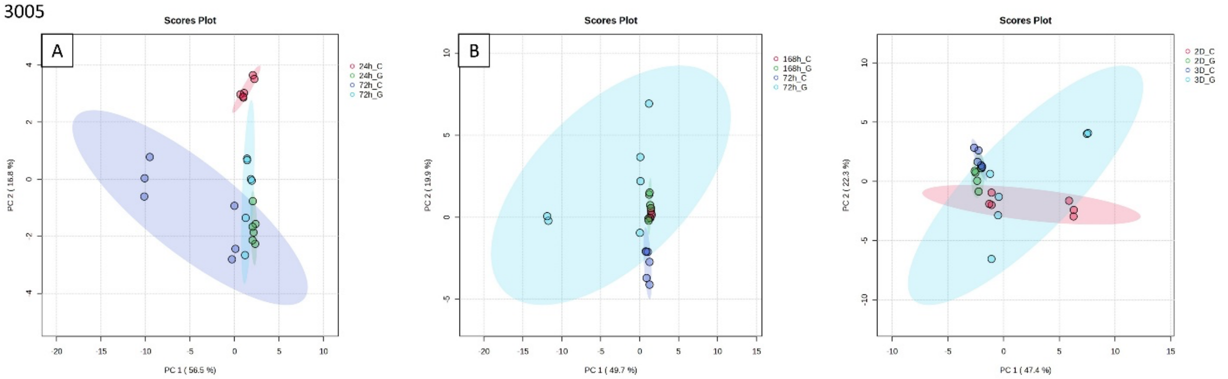

3019

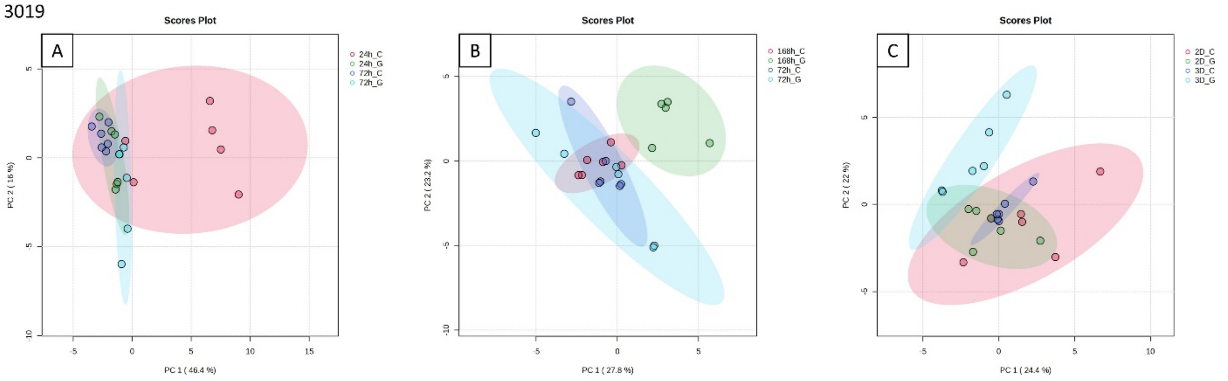

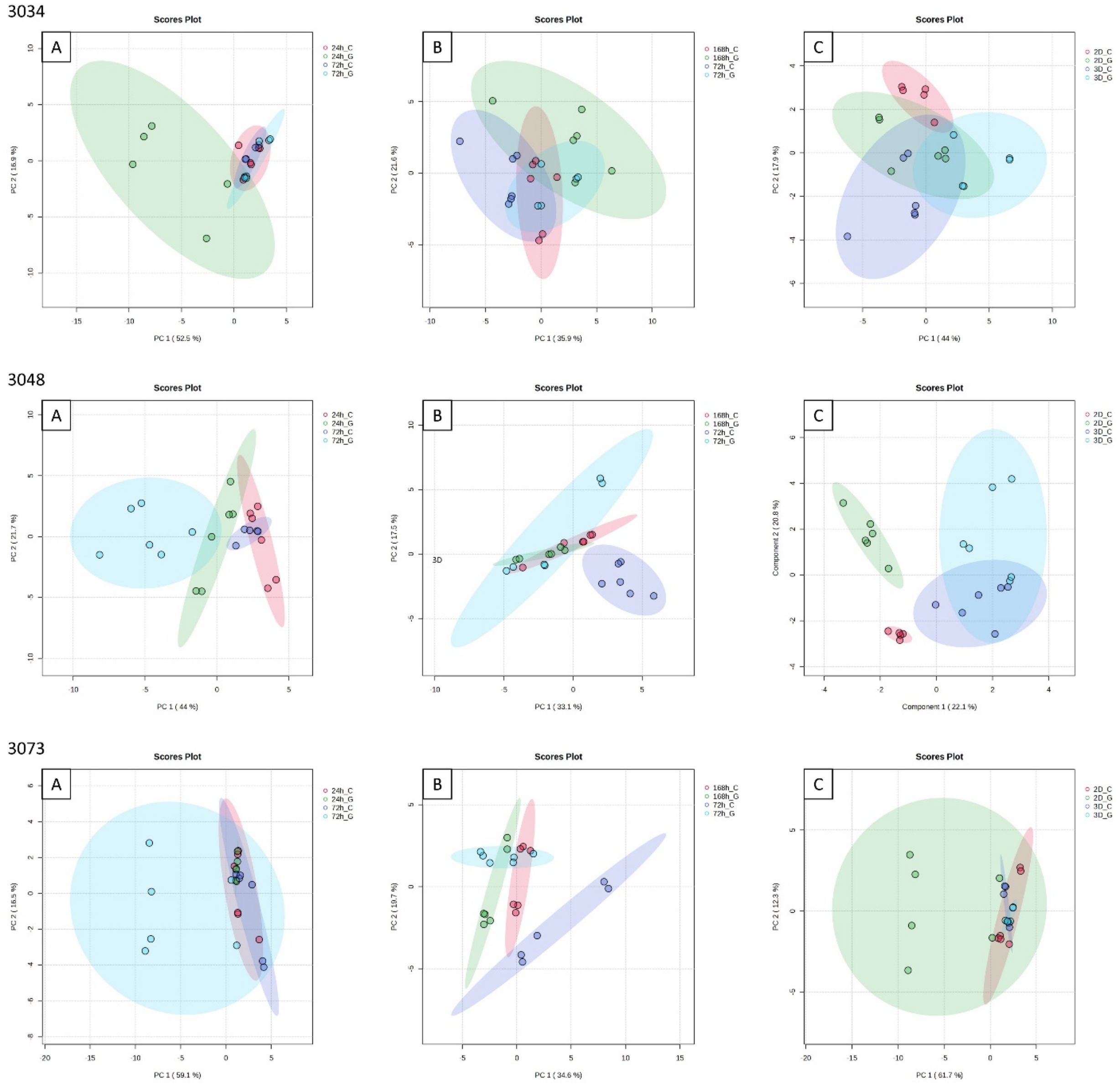

Figure S2. PCA score plots showing separation of all cell lines in A) 2D 24h vs 72h, B) 3D 72h vs 168h, C) 72h 2D vs 3D (n=6).

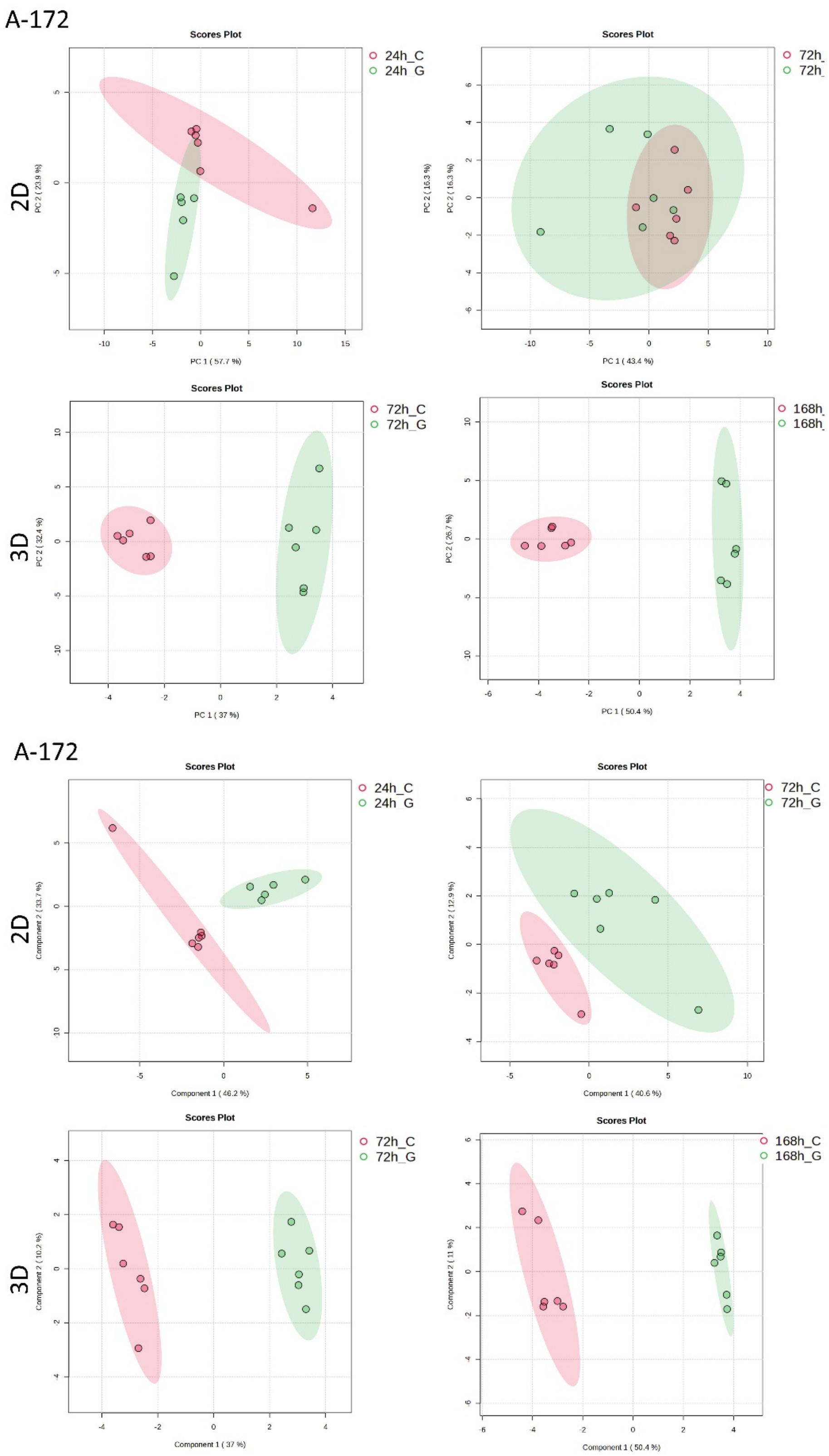

Figutre S3. PCA (top) and PLS-DA scrore plots showing separations of A-172 cell line in 2D and 3D, treated (G) and untreated (C).

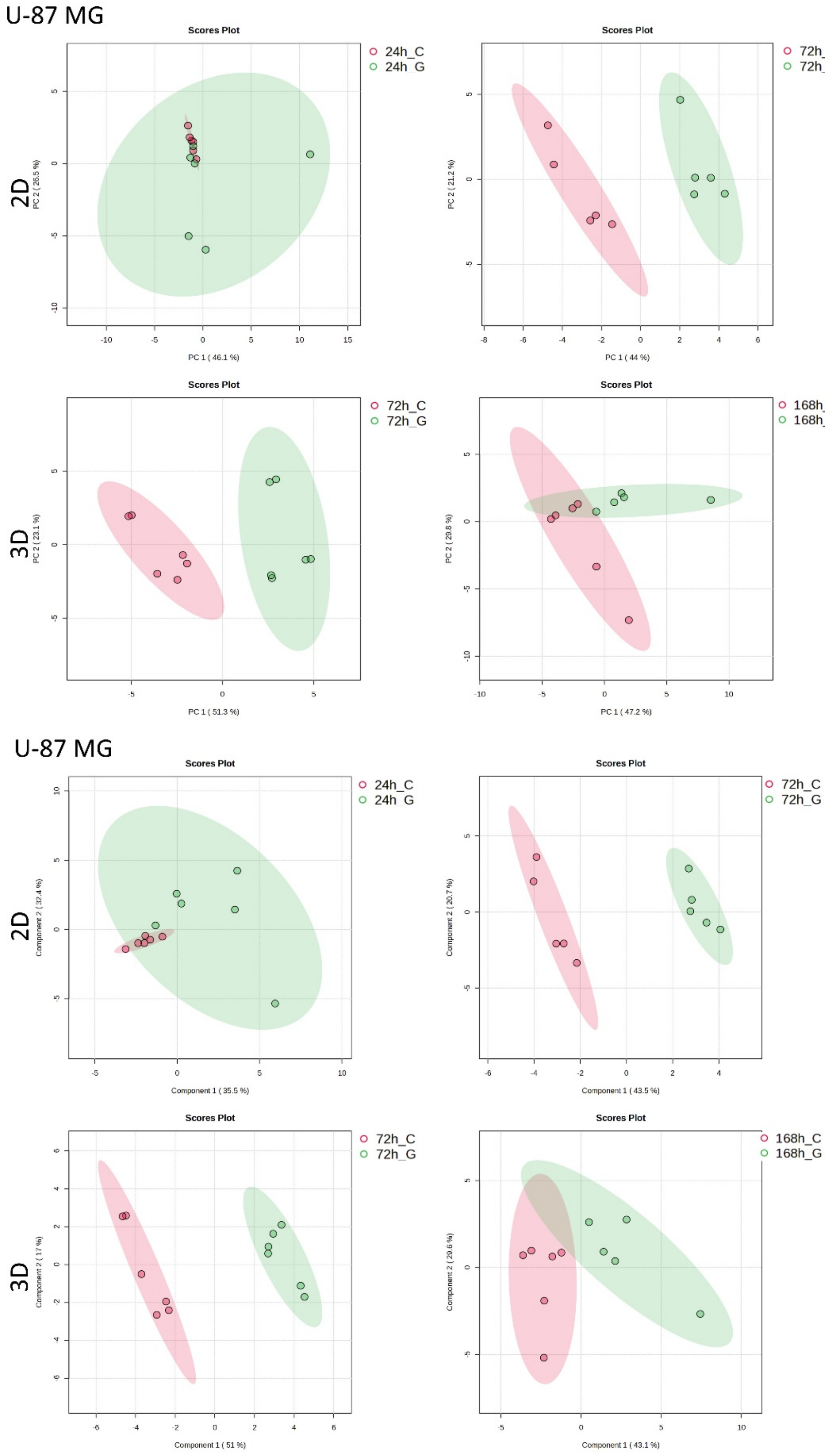

Figutre S4. PCA (top) and PLS-DA scrore plots showing separations of U-87 MG cell line in 2D and 3D, treated (G) and untreated (C).

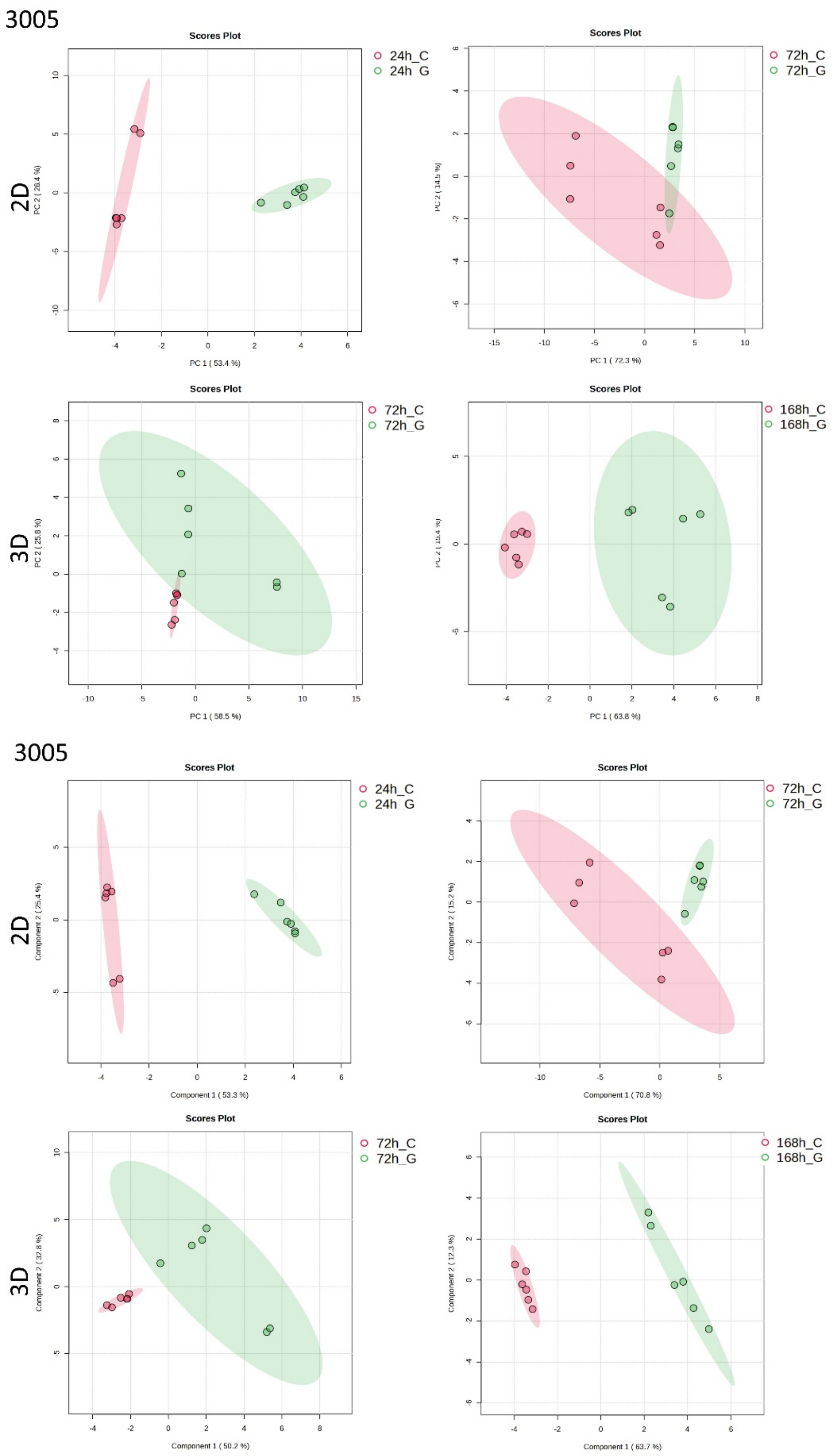

Figutre S5. PCA (top) and PLS-DA scrore plots showing separations of 3005 MG cell line in 2D and 3D, treated (G) and untreated (C).

3019

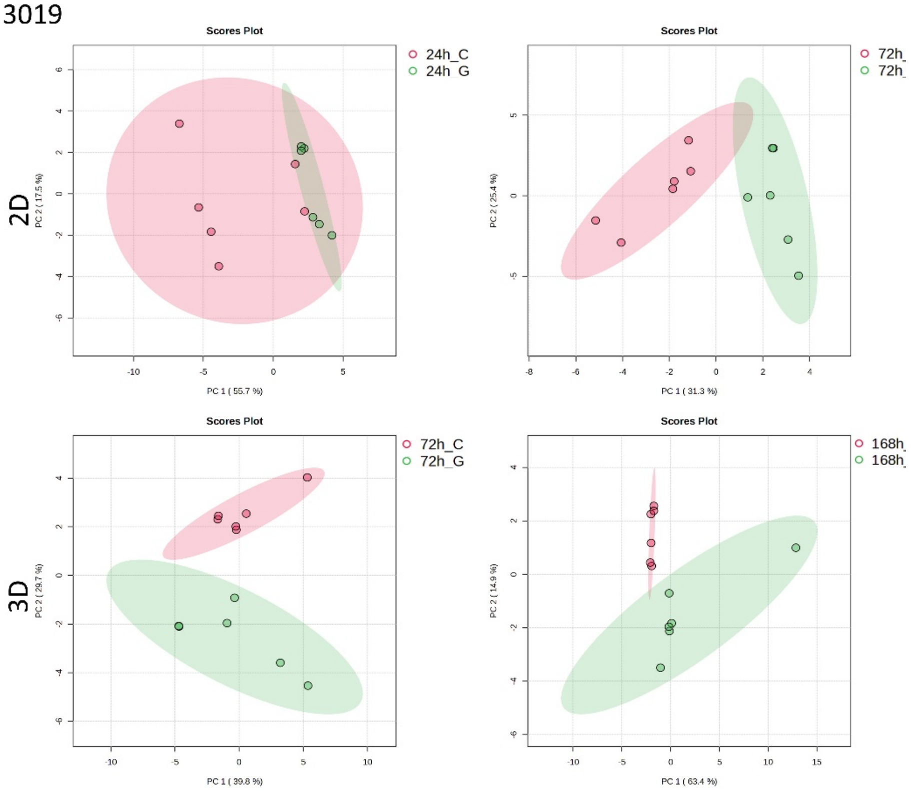

3019

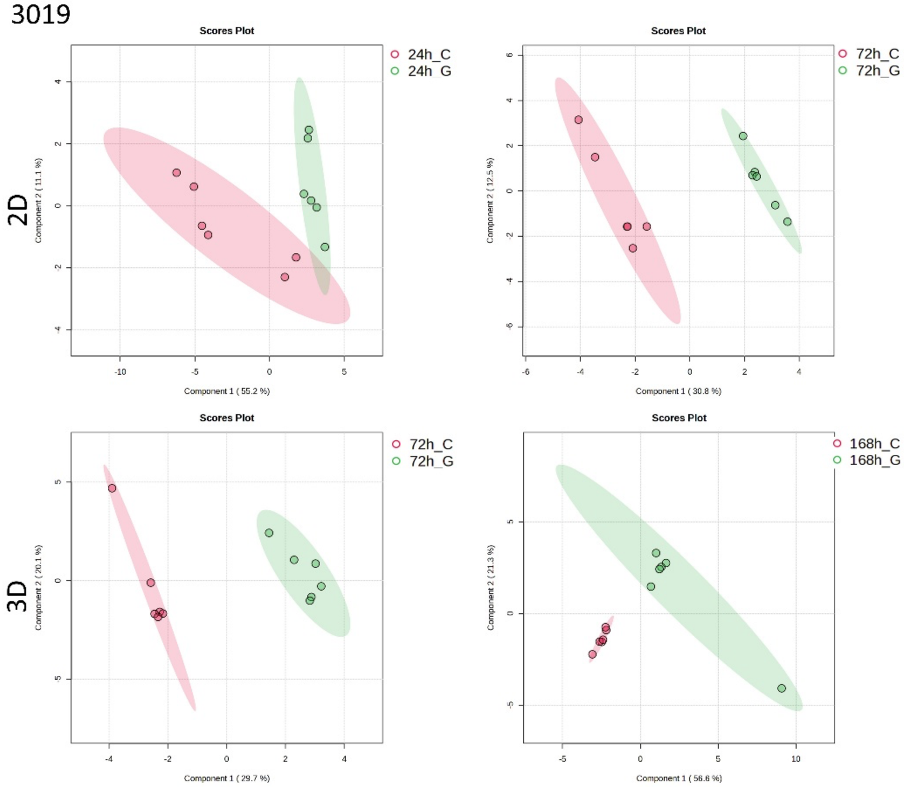

Figutre S6. PCA (top) and PLS-DA scrore plots showing separations of 3019 MG cell line in 2D and 3D, treated (G) and untreated (C).

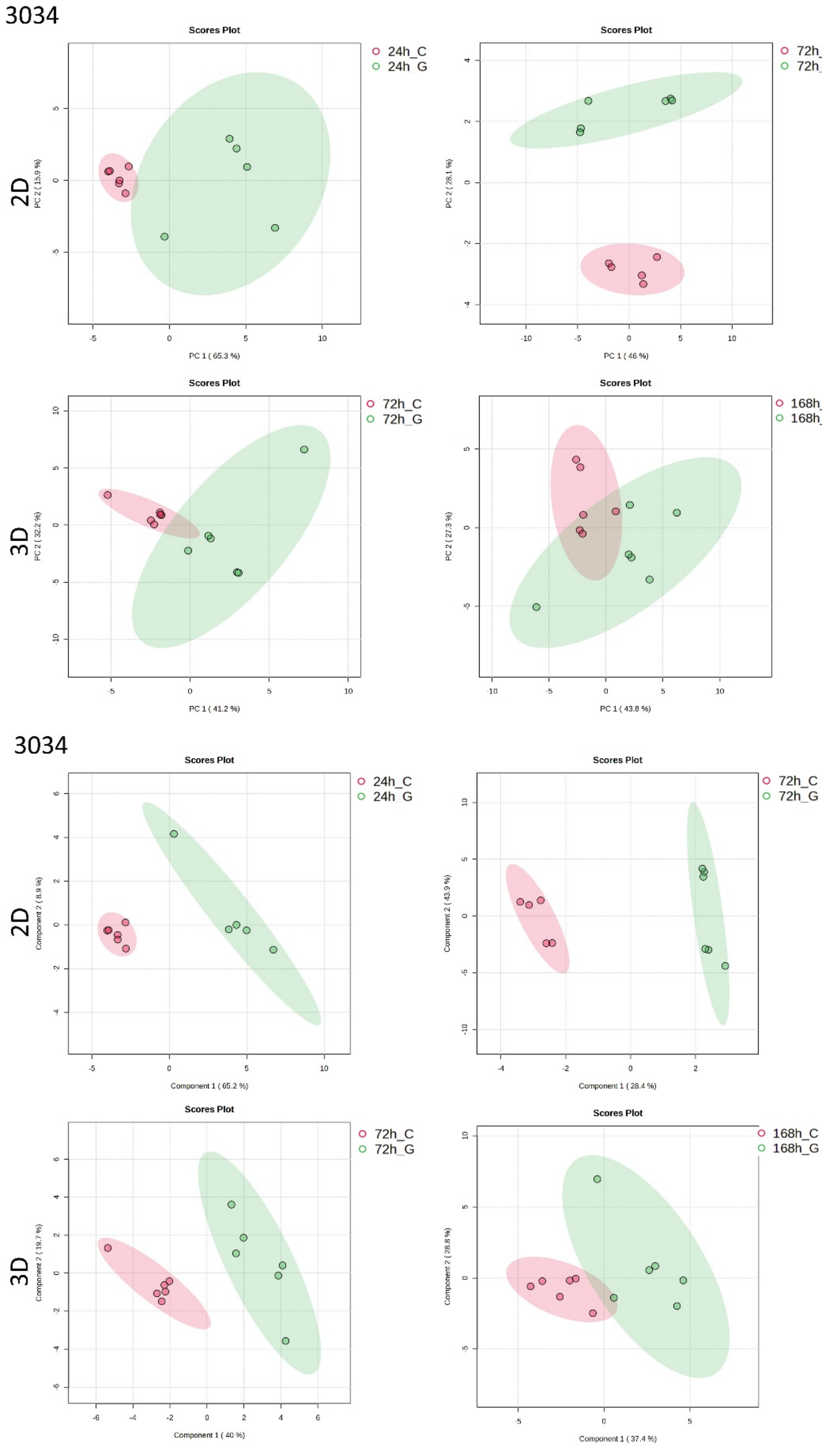

Figutre S7. PCA (top) and PLS-DA scrore plots showing separations of 3034 MG cell line in 2D and 3D, treated (G) and untreated (C).

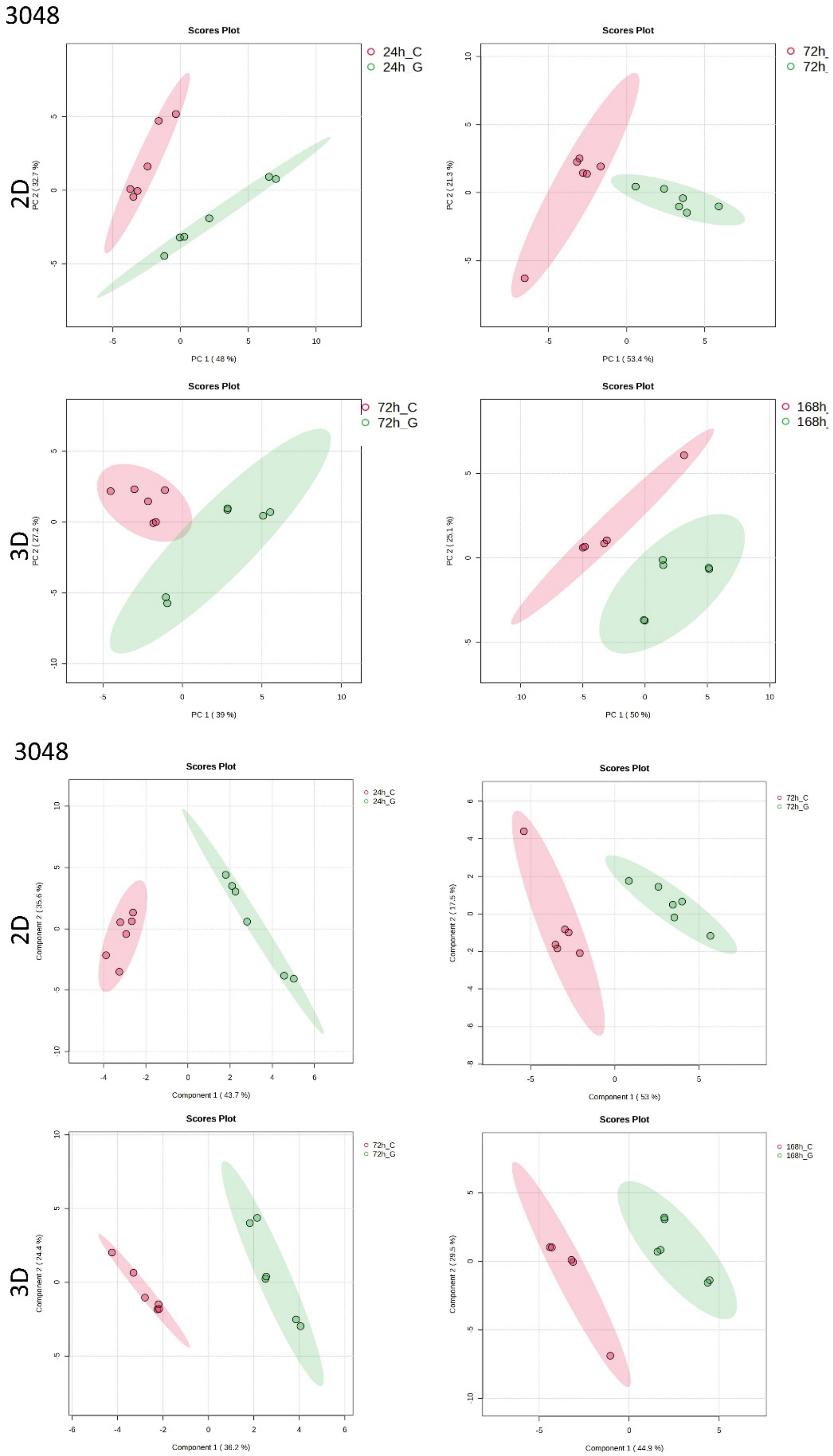

Figutre S8. PCA (top) and PLS-DA scrore plots showing separations of 3048 MG cell line in 2D and 3D, treated (G) and untreated (C).

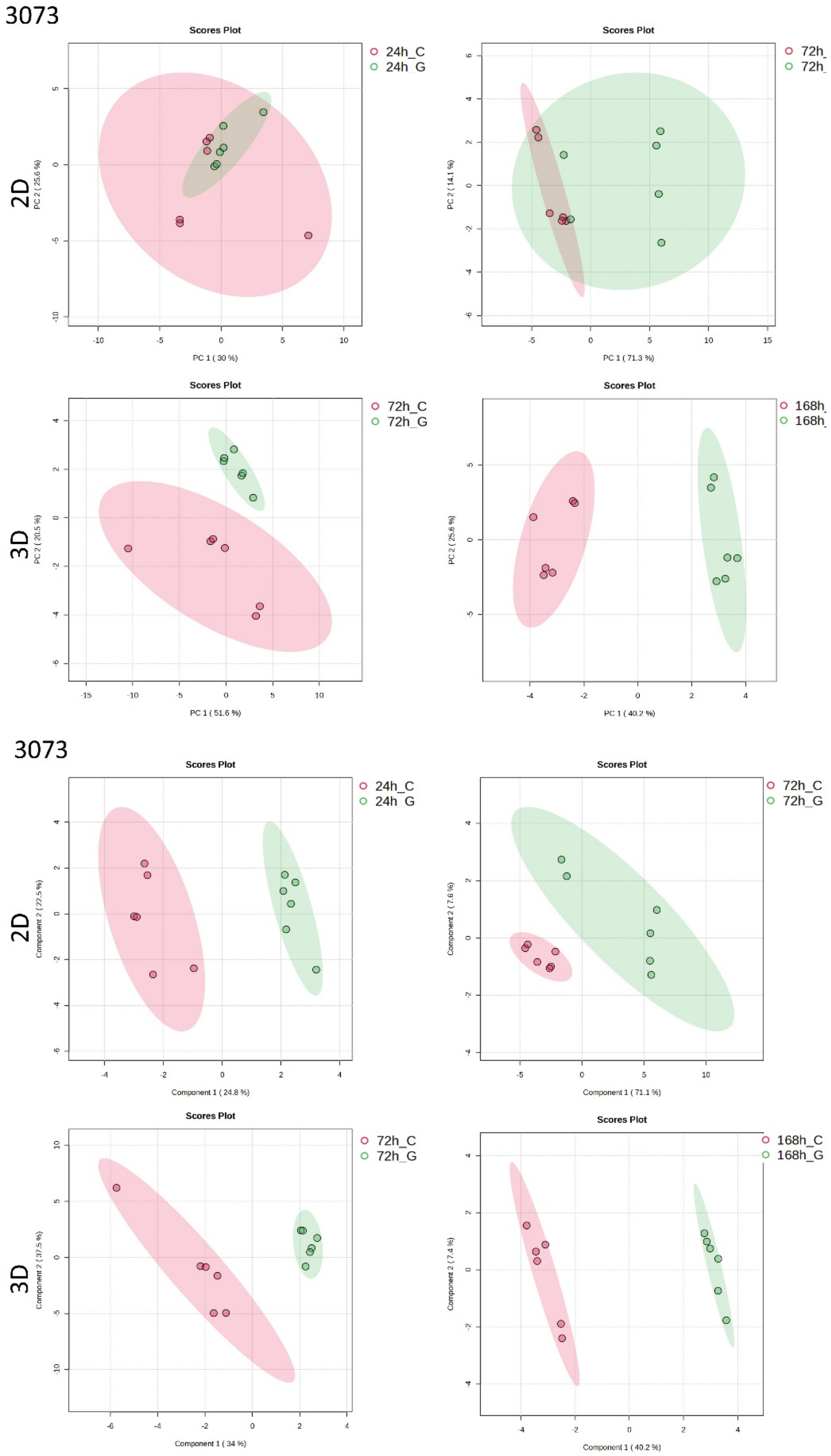

Figutre S9. PCA (top) and PLS-DA scrore plots showing separations of 3073 MG cell line in 2D and 3D, treated (G) and untreated (C).

Table S2. Cross-validated PLS-DA performance across GBM models.

| Cell Line | Format | Time | Comparison | Best Components | Accuracy | Q2 | R2 |
|---|---|---|---|---|---|---|---|
| 3005 | 2D | 24h_72h | 2D_24h_72h | 4 | 0.96 | 0.65474 | 0.84238 |
| | 2D | 24h | 2D_24h | 3 | 1 | 0.97996 | 0.99671 |
| | 2D_3D | 72h | 2D_3D_72h | 3 | 0.70667 | 0.65796 | 0.88375 |
| | 2D | 72h | 2D_72h | 4 | 1 | 0.89178 | 0.98585 |
| | 3D | 168h | 3D_168h | 4 | 1 | 0.98378 | 0.99883 |
| | 3D | 72h_168h | 3D_72h_168h | 3 | 0.79333 | 0.68473 | 0.90763 |
| | 3D | 72h | 3D_72h | 6 | 0.95 | 0.89893 | 0.99977 |
| 3019 | 2D | 24h_72h | 2D_24h_72h | 2 | 0.62 | 0.35086 | 0.57689 |
| | 2D | 24h | 2D_24h | 4 | 0.8 | 0.69851 | 0.94982 |
| | 2D_3D | 72h | 2D_3D_72h | 3 | 0.77143 | 0.87795 | 0.92452 |
| | 2D | 72h | 2D_72h | 3 | 1 | 0.97771 | 0.99513 |
| | 3D | 168h | 3D_168h | 2 | 1 | 0.7575 | 0.96044 |
| | 3D | 72h_168h | 3D_72h_168h | 2 | 0.47333 | 0.19839 | 0.53053 |
| | 3D | 72h | 3D_72h | 5 | 1 | 0.97565 | 0.99942 |
| 3034 | 2D | 24h_72h | 2D_24h_72h | 3 | 0.82 | 0.91722 | 0.96921 |
| | 2D | 24h | 2D_24h | 5 | 1 | 0.92275 | 0.99257 |
| | 2D_3D | 72h | 2D_3D_72h | 4 | 0.82 | 0.82507 | 0.94573 |
| | 2D | 72h | 2D_72h | 2 | 0.83333 | -0.30939 | 0.94595 |
| | 3D | 168h | 3D_168h | 1 | 0.9 | 0.39949 | 0.71562 |
| | 3D | 72h_168h | 3D_72h_168h | 2 | 0.475 | 0.76311 | 0.87437 |
| | 3D | 72h | 3D_72h | 3 | 1 | 0.90148 | 0.97777 |
| 3048 | 2D | 24h_72h | 2D_24h_72h | 3 | 0.73333 | 0.74137 | 0.86342 |
| | 2D | 24h | 2D_24h | 3 | 1 | 0.95536 | 0.99102 |
| | 2D_3D | 72h | 2D_3D_72h | 4 | 0.93333 | 0.91498 | 0.96714 |
| | 2D | 72h | 2D_72h | 4 | 1 | 0.95217 | 0.99529 |
| | 3D | 168h | 3D_168h | 5 | 1 | 0.98214 | 0.99866 |
| | 3D | 72h_168h | 3D_72h_168h | 2 | 0.56333 | 0.57985 | 0.86422 |
| | 3D | 72h | 3D_72h | 5 | 1 | 0.98625 | 0.99899 |
| 3073 | 2D | 24h_72h | 2D_24h_72h | 5 | 0.83 | 0.80533 | 0.94726 |
| | 2D | 24h | 2D_24h | 5 | 0.93333 | 0.81003 | 0.99811 |
| | 2D_3D | 72h | 2D_3D_72h | 4 | 0.82 | 0.57748 | 0.85255 |
| | 2D | 72h | 2D_72h | 5 | 1 | 0.93482 | 0.99616 |
| | 3D | 168h | 3D_168h | 4 | 1 | 0.98378 | 0.99883 |
| | 3D | 72h_168h | 3D_72h_168h | 3 | 0.88 | 0.65929 | 0.87341 |
| | 3D | 72h | 3D_72h | 5 | 1 | 0.96631 | 0.99908 |
| A-172 | 2D | 24h | 2D_24h | 5 | 1 | 0.93118 | 0.99647 |

| | | | | | | | |
|---|---|---|---|---|---|---|---|
| | 2D_3D | 72h | 2D_3D_72h | 3 | 0.64 | 0.49049 | 0.87458 |
| | 2D | 72h | 2D_72h | 4 | 0.8 | 0.58358 | 0.95365 |
| | 3D | 168h | 3D_168h | 2 | 1 | 0.98727 | 0.99521 |
| | 3D | 72h+168h | 3D_72h+168h | 4 | 1 | 0.95995 | 0.99722 |
| | 3D | 72h_168h | 3D_72h_168h | 5 | 0.96 | 0.92071 | 0.97342 |
| | 2D | 24h_72h | 2D_24h_72h | 4 | 0.86 | 0.76004 | 0.9672 |
| U-87 MG | 2D | 24h_72h | 2D_24h_72h | 4 | 0.83 | 0.68205 | 0.93221 |
| | 2D | 24h | 2D_24h | 1 | 0.95 | -0.35231 | 0.75069 |
| | 2D_3D | 72h | 2D_3D_72h | 5 | 1 | 0.74933 | 0.98245 |
| | 2D | 72h | 2D_72h | 3 | 0.83333 | 0.40482 | 0.84569 |
| | 3D | 168h | 3D_168h | 1 | 0.9 | 0.39167 | 0.69546 |
| | 3D | 72h_168h | 3D_72h_168h | 2 | 0.78 | 0.91827 | 0.95246 |
| | 3D | 72h | 3D_72h | 5 | 1 | 0.98934 | 0.99951 |

Table S3. Comprehensive metabolite panel by cell line (3005, 3019, 3034, 3048, 3073, A-172, U-87 MG): VIP, FDR, stars

| Cell Line | Format | Time | Condition | Metabolite | VIP | FDR | stars |
|---|---|---|---|---|---|---|---|
| 3005 | 2D | 24h | 2D_24h | Allantoin | 1.2467 | 0.008434 | ** |
| | 3D | 168h | 3D_168h | Allantoin | 1.0036 | 0.006815 | ** |
| | 2D | 24h | 2D_24h | Choline | 1.3072 | 0.008434 | ** |
| | 3D | 168h | 3D_168h | Choline | 1.1882 | 0.003968 | ** |
| | 2D | 24h | 2D_24h | Cysteine | 1.0935 | 0.006494 | ** |
| | 2D | 72h | 2D_72h | Cysteine | 1.299 | 0.011688 | * |
| | 3D | 168h | 3D_168h | Cysteine | 1.1649 | 0.003968 | ** |
| | 3D | 72h | 3D_72h | Cysteine | 1.5496 | 0.02381 | * |
| | 2D | 24h | 2D_24h | Glutamicacid | 1.0655 | 0.006494 | ** |
| | 2D | 72h | 2D_72h | Glutamine | 1.3858 | 0.011688 | * |
| | 3D | 168h | 3D_168h | Glutamine | 1.254 | 0.003968 | ** |
| | 3D | 72h | 3D_72h | Glutamine | 1.5778 | 0.015873 | * |
| | 3D | 168h | 3D_168h | Histidine | 1.2612 | 0.003968 | ** |
| | 2D | 72h | 2D_72h | Isoleucine | 1.2834 | 0.011688 | * |
| | 2D | 24h | 2D_24h | Lysine | 1.2104 | 0.006494 | ** |
| | 3D | 168h | 3D_168h | Lysine | 1.1372 | 0.003968 | ** |
| | 2D | 72h | 2D_72h | Methionine | 1.1695 | 0.02904 | * |
| | 3D | 168h | 3D_168h | Methionine | 1.09 | 0.003968 | ** |
| | 2D | 24h | 2D_24h | Niacinamide | 1.1203 | 0.008434 | ** |
| | 2D | 72h | 2D_72h | Niacinamide | 1.3112 | 0.014994 | * |
| | 3D | 168h | 3D_168h | Niacinamide | 1.0173 | 0.003968 | ** |
| | 3D | 72h | 3D_72h | Niacinamide | 1.6055 | 0.015873 | * |
| | 3D | 72h | 3D_72h | Ornithine | 1.4831 | 0.031746 | * |
| | 2D | 24h | 2D_24h | Phenylalanine | 1.1686 | 0.006494 | ** |
| | 2D | 72h | 2D_72h | Phenylalanine | 1.2333 | 0.011688 | * |
| | 3D | 168h | 3D_168h | Phenylalanine | 1.2398 | 0.003968 | ** |
| | 2D | 24h | 2D_24h | Proline | 1.3449 | 0.006494 | ** |
| | 3D | 168h | 3D_168h | Proline | 1.1253 | 0.003968 | ** |
| | 2D | 72h | 2D_72h | Serine | 1.126 | 0.025213 | * |
| | 3D | 168h | 3D_168h | Serine | 1.0912 | 0.003968 | ** |
| | 2D | 24h | 2D_24h | Threonine | 1.0547 | 0.013751 | * |
| | 2D | 24h | 2D_24h | Tryptophan | 1.258 | 0.008434 | ** |
| | 2D | 72h | 2D_72h | Tryptophan | 1.2273 | 0.014994 | * |
| | 3D | 168h | 3D_168h | Tryptophan | 1.0225 | 0.006815 | ** |
| | 2D | 24h | 2D_24h | Tyrosine | 1.2133 | 0.006494 | ** |
| | 2D | 72h | 2D_72h | Tyrosine | 1.0667 | 0.023377 | * |
| | 2D | 24h | 2D_24h | Uracil | 1.2965 | 0.008434 | ** |
| | 2D | 72h | 2D_72h | Uracil | 1.3756 | 0.011688 | * |
| | 3D | 168h | 3D_168h | Uracil | 1.2685 | 0.003968 | ** |

| | | | | | | | |
|---|---|---|---|---|---|---|---|
| | 3D | 72h | 3D_72h | Uracil | 1.4161 | 0.015873 | * |
| | 2D | 24h | 2D_24h | Valine | 1.2714 | 0.006494 | ** |
| 3019 | 2D | 72h | 2D_72h | Allantoin | 1.1145 | 0.019481 | * |
| | 3D | 72h | 3D_72h | Allantoin | 1.0505 | 0.01461 | * |
| | 3D | 72h | 3D_72h | Glutamicacid | 1.3095 | 0.019481 | * |
| | 3D | 72h | 3D_72h | Histidine | 1.5642 | 0.01461 | * |
| | 2D | 24h | 2D_24h | Lysine | 1.3021 | 0.040909 | * |
| | 3D | 168h | 3D_168h | Lysine | 1.0544 | 0.007305 | ** |
| | 2D | 24h | 2D_24h | Methionine | 1.4629 | 0.011688 | * |
| | 3D | 168h | 3D_168h | Methionine | 1.5066 | 0.007305 | ** |
| | 2D | 24h | 2D_24h | Niacinamide | 1.3656 | 0.027053 | * |
| | 2D | 24h | 2D_24h | Ornithine | 1.3793 | 0.040909 | * |
| | 2D | 24h | 2D_24h | Proline | 1.4157 | 0.019278 | * |
| | 3D | 168h | 3D_168h | Serine | 1.0485 | 0.007305 | ** |
| | 2D | 72h | 2D_72h | Tryptophan | 1.631 | 0.02699 | * |
| | 2D | 24h | 2D_24h | Tyrosine | 1.3967 | 0.011688 | * |
| | 3D | 168h | 3D_168h | Tyrosine | 1.4581 | 0.007305 | ** |
| | 2D | 24h | 2D_24h | Uracil | 1.5132 | 0.011688 | * |
| | 2D | 72h | 2D_72h | Uracil | 1.8454 | 0.02699 | * |
| | 3D | 72h | 3D_72h | Uracil | 1.8903 | 0.01461 | * |
| | 2D | 24h | 2D_24h | Valine | 1.3542 | 0.011688 | * |
| 3034 | 2D | 24h | 2D_24h | Adenosine | 1.1594 | 0.011688 | * |
| | 3D | 72h | 3D_72h | Allantoin | 1.6015 | 0.00974 | ** |
| | 2D | 24h | 2D_24h | Glutamicacid | 1.1628 | 0.011688 | * |
| | 3D | 72h | 3D_72h | Glutamine | 1.4951 | 0.00974 | ** |
| | 2D | 24h | 2D_24h | Histidine | 1.3424 | 0.011688 | * |
| | 2D | 24h | 2D_24h | Leucine | 1.2506 | 0.011688 | * |
| | 2D | 24h | 2D_24h | Lysine | 1.191 | 0.011688 | * |
| | 3D | 72h | 3D_72h | Methionine | 1.5239 | 0.00974 | ** |
| | 3D | 72h | 3D_72h | Niacinamide | 1.4326 | 0.00974 | ** |
| | 2D | 24h | 2D_24h | Proline | 1.2001 | 0.019481 | * |
| | 2D | 24h | 2D_24h | Tryptophan | 1.2562 | 0.011688 | * |
| | 3D | 168h | 3D_168h | Tryptophan | 1.3983 | 0.044983 | * |
| | 3D | 72h | 3D_72h | Tryptophan | 1.2669 | 0.033395 | * |
| | 3D | 168h | 3D_168h | Tyrosine | 1.6383 | 0.044983 | * |
| | 2D | 24h | 2D_24h | Uracil | 1.2789 | 0.011688 | * |
| | 2D | 72h | 2D_72h | Uracil | 1.8786 | 0.029221 | * |
| | 3D | 72h | 3D_72h | Uracil | 1.6662 | 0.00974 | ** |
| | 2D | 24h | 2D_24h | Valine | 1.1976 | 0.011688 | * |
| 3048 | 2D | 24h | 2D_24h | Allantoin | 1.3721 | 0.007305 | ** |
| | 2D | 72h | 2D_72h | Allantoin | 1.2524 | 0.008349 | ** |
| | 3D | 168h | 3D_168h | Allantoin | 1.6831 | 0.015004 | * |
| | 2D | 24h | 2D_24h | Cysteine | 1.0207 | 0.007305 | ** |

| | | | | | | | |
|---|---|---|---|---|---|---|---|
| | 2D | 72h | 2D_72h | Glutamicacid | 1.1458 | 0.005313 | ** |
| | 2D | 72h | 2D_72h | Glutamine | 1.3019 | 0.008245 | ** |
| | 3D | 72h | 3D_72h | Glutamine | 1.5045 | 0.00974 | ** |
| | 2D | 24h | 2D_24h | Histidine | 1.2581 | 0.007305 | ** |
| | 2D | 24h | 2D_24h | Isoleucine | 1.1973 | 0.007305 | ** |
| | 2D | 72h | 2D_72h | Isoleucine | 1.2135 | 0.008349 | ** |
| | 2D | 24h | 2D_24h | Leucine | 1.1596 | 0.023377 | * |
| | 2D | 72h | 2D_72h | Leucine | 1.0581 | 0.005313 | ** |
| | 3D | 168h | 3D_168h | Leucine | 1.3655 | 0.033395 | * |
| | 2D | 72h | 2D_72h | Lysine | 1.3763 | 0.005313 | ** |
| | 2D | 72h | 2D_72h | Methionine | 1.4176 | 0.005313 | ** |
| | 3D | 168h | 3D_168h | Methionine | 1.4213 | 0.01461 | * |
| | 3D | 72h | 3D_72h | Niacinamide | 1.4832 | 0.00974 | ** |
| | 3D | 168h | 3D_168h | Ornithine | 1.2558 | 0.033395 | * |
| | 2D | 72h | 2D_72h | Proline | 1.237 | 0.005313 | ** |
| | 3D | 72h | 3D_72h | Proline | 1.211 | 0.045455 | * |
| | 2D | 24h | 2D_24h | Serine | 1.381 | 0.007305 | ** |
| | 2D | 72h | 2D_72h | Serine | 1.2242 | 0.005313 | ** |
| | 2D | 72h | 2D_72h | Tryptophan | 1.3436 | 0.005313 | ** |
| | 3D | 168h | 3D_168h | Tryptophan | 1.2142 | 0.01461 | * |
| | 3D | 72h | 3D_72h | Tryptophan | 1.3644 | 0.00974 | ** |
| | 2D | 24h | 2D_24h | Tyrosine | 1.3969 | 0.007305 | ** |
| | 3D | 72h | 3D_72h | Tyrosine | 1.273 | 0.00974 | ** |
| | 2D | 72h | 2D_72h | Uracil | 1.2202 | 0.005313 | ** |
| | 3D | 168h | 3D_168h | Uracil | 1.4293 | 0.01461 | * |
| | 3D | 72h | 3D_72h | Uracil | 1.6387 | 0.00974 | ** |
| | 2D | 24h | 2D_24h | Valine | 1.3179 | 0.007305 | ** |
| | 3D | 168h | 3D_168h | Valine | 1.4871 | 0.01461 | * |
| | 3D | 72h | 3D_72h | Valine | 1.2804 | 0.045455 | * |
| 3073 | 3D | 168h | 3D_168h | Adenosine | 1.0458 | 0.046753 | * |
| | 3D | 168h | 3D_168h | Allantoin | 1.1043 | 0.034091 | * |
| | 3D | 72h | 3D_72h | Allantoin | 1.6839 | 0.03221 | * |
| | 2D | 72h | 2D_72h | Cysteine | 1.1962 | 0.021251 | * |
| | 3D | 168h | 3D_168h | Cysteine | 1.4669 | 0.008349 | ** |
| | 2D | 24h | 2D_24h | Glutamine | 1.4927 | 0.029221 | * |
| | 2D | 72h | 2D_72h | Glutamine | 1.1767 | 0.021251 | * |
| | 3D | 168h | 3D_168h | Histidine | 1.5285 | 0.008349 | ** |
| | 2D | 72h | 2D_72h | Isoleucine | 1.2023 | 0.008349 | ** |
| | 3D | 168h | 3D_168h | Isoleucine | 1.1648 | 0.012987 | * |
| | 3D | 168h | 3D_168h | Lysine | 1.412 | 0.008349 | ** |
| | 2D | 72h | 2D_72h | Methionine | 1.1904 | 0.008349 | ** |
| | 3D | 72h | 3D_72h | Methionine | 1.3647 | 0.046753 | * |
| | 2D | 72h | 2D_72h | Niacinamide | 1.3236 | 0.008349 | ** |

| | | | | | | | |
|---|---|---|---|---|---|---|---|
| | 2D | 72h | 2D_72h | Phenylalanine | 1.1759 | 0.008349 | ** |
| | 3D | 168h | 3D_168h | Phenylalanine | 1.5349 | 0.008349 | ** |
| | 2D | 24h | 2D_24h | Proline | 1.8999 | 0.029221 | * |
| | 3D | 168h | 3D_168h | Proline | 1.0172 | 0.046753 | * |
| | 2D | 72h | 2D_72h | Serine | 1.2129 | 0.01461 | * |
| | 3D | 168h | 3D_168h | Serine | 1.3256 | 0.008349 | ** |
| | 2D | 72h | 2D_72h | Threonine | 1.188 | 0.021251 | * |
| | 3D | 168h | 3D_168h | Threonine | 1.2329 | 0.008349 | ** |
| | 2D | 72h | 2D_72h | Tryptophan | 1.2397 | 0.008349 | ** |
| | 3D | 168h | 3D_168h | Tryptophan | 1.1327 | 0.046753 | * |
| | 3D | 72h | 3D_72h | Tryptophan | 1.688 | 0.029221 | * |
| | 2D | 72h | 2D_72h | Uracil | 1.4124 | 0.008349 | ** |
| | 3D | 168h | 3D_168h | Uracil | 1.2753 | 0.023377 | * |
| | 3D | 72h | 3D_72h | Uracil | 1.7576 | 0.029221 | * |
| | 2D | 24h | 2D_24h | Valine | 1.6083 | 0.029221 | * |
| | 3D | 168h | 3D_168h | Valine | 1.2935 | 0.012987 | * |
| | 3D | 72h | 3D_72h | Valine | 1.6379 | 0.03221 | * |
| A-172 | 2D | 24h | 2D_24h | Allantoin | 1.8205 | 0.012987 | * |
| | 3D | 168h | 3D_168h | Allantoin | 1.4139 | 0.006251 | ** |
| | 3D | 72h | 3D_72h | Allantoin | 1.6185 | 0.008349 | ** |
| | 3D | 168h | 3D_168h | Cysteine | 1.2199 | 0.005313 | ** |
| | 3D | 72h | 3D_72h | Cysteine | 1.2619 | 0.008349 | ** |
| | 3D | 168h | 3D_168h | Glutamine | 1.3853 | 0.005313 | ** |
| | 3D | 168h | 3D_168h | Histidine | 1.1009 | 0.005313 | ** |
| | 3D | 168h | 3D_168h | Leucine | 1.2151 | 0.005313 | ** |
| | 3D | 168h | 3D_168h | Lysine | 1.3021 | 0.005313 | ** |
| | 3D | 72h | 3D_72h | Lysine | 1.4465 | 0.008349 | ** |
| | 3D | 168h | 3D_168h | Methionine | 1.2729 | 0.005313 | ** |
| | 3D | 72h | 3D_72h | Methionine | 1.3674 | 0.008349 | ** |
| | 3D | 168h | 3D_168h | Niacinamide | 1.0925 | 0.005313 | ** |
| | 2D | 24h | 2D_24h | Ornithine | 1.4682 | 0.012987 | * |
| | 3D | 168h | 3D_168h | Ornithine | 1.0044 | 0.016698 | * |
| | 3D | 72h | 3D_72h | Ornithine | 1.4309 | 0.008349 | ** |
| | 3D | 168h | 3D_168h | Phenylalanine | 1.3361 | 0.005313 | ** |
| | 3D | 168h | 3D_168h | Serine | 1.266 | 0.005313 | ** |
| | 3D | 72h | 3D_72h | Serine | 1.2128 | 0.008349 | ** |
| | 2D | 24h | 2D_24h | Tryptophan | 1.6864 | 0.012987 | * |
| | 3D | 168h | 3D_168h | Tyrosine | 1.1438 | 0.008991 | ** |
| | 2D | 24h | 2D_24h | Uracil | 1.6579 | 0.012987 | * |
| | 3D | 168h | 3D_168h | Uracil | 1.4003 | 0.005313 | ** |
| | 3D | 72h | 3D_72h | Uracil | 1.6082 | 0.008349 | ** |
| U87MG | 3D | 72h | 3D_72h | Adenosine | 1.2438 | 0.005313 | ** |
| | 3D | 72h | 3D_72h | Allantoin | 1.4314 | 0.006251 | ** |

| | | | | | | | |
|---|---|---|---|---|---|---|---|
| | 3D | 72h | 3D_72h | Choline | 1.1484 | 0.005313 | ** |
| | 3D | 72h | 3D_72h | Glutamicacid | 1.2518 | 0.005313 | ** |
| | 3D | 72h | 3D_72h | Histidine | 1.1785 | 0.005313 | ** |
| | 3D | 168h | 3D_168h | Methionine | 1.1634 | 0.046753 | * |
| | 3D | 72h | 3D_72h | Methionine | 1.3087 | 0.005313 | ** |
| | 3D | 168h | 3D_168h | Niacinamide | 1.5357 | 0.029221 | * |
| | 3D | 168h | 3D_168h | Proline | 1.5211 | 0.029221 | * |
| | 3D | 72h | 3D_72h | Serine | 1.1842 | 0.005313 | ** |
| | 3D | 72h | 3D_72h | Tryptophan | 1.1085 | 0.005313 | ** |
| | 3D | 72h | 3D_72h | Tyrosine | 1.0347 | 0.027273 | * |
| | 3D | 168h | 3D_168h | Uracil | 1.24 | 0.029221 | * |
| | 3D | 72h | 3D_72h | Uracil | 1.4291 | 0.005313 | ** |
| | 3D | 168h | 3D_168h | Valine | 1.3233 | 0.029221 | * |

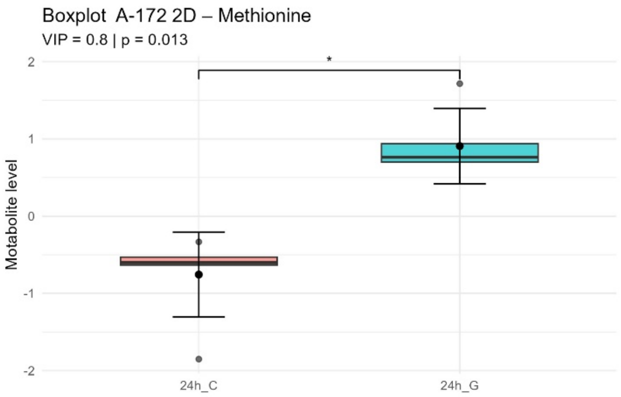
Boxplot A-172 2D – Methionine
VIP = 0.8 | p = 0.013
*
Motabolite level
2
1
0
-1
-2
24h_C
24h_G

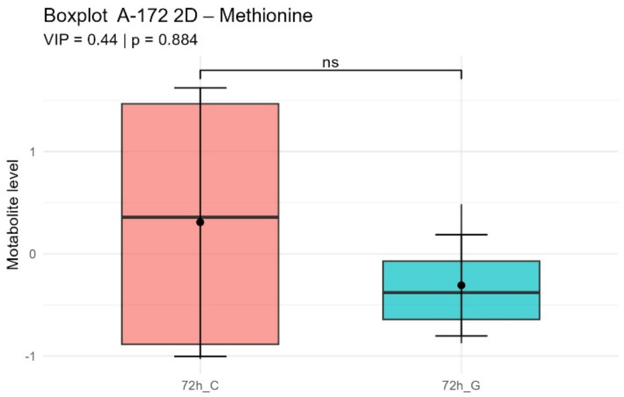
Boxplot A-172 2D – Methionine
VIP = 0.44 | p = 0.884
ns
Motabolite level
1
0
-1
72h_C
72h_G

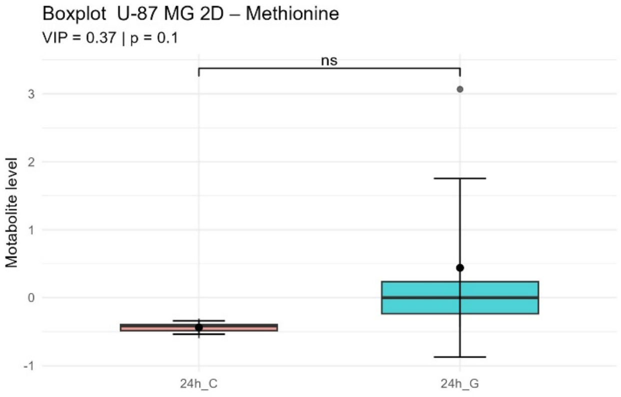
Boxplot U-87 MG 2D – Methionine
VIP = 0.37 | p = 0.1
ns
Motabolite level
3
2
1
0
-1
24h_C
24h_G

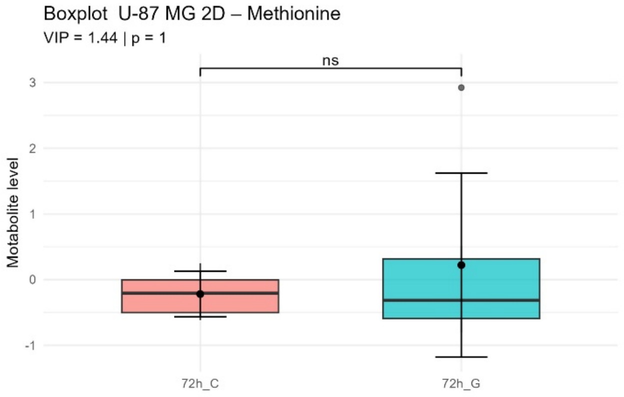
Boxplot U-87 MG 2D – Methionine
VIP = 1.44 | p = 1
ns
Motabolite level
3
2
1
0
-1
72h_C
72h_G

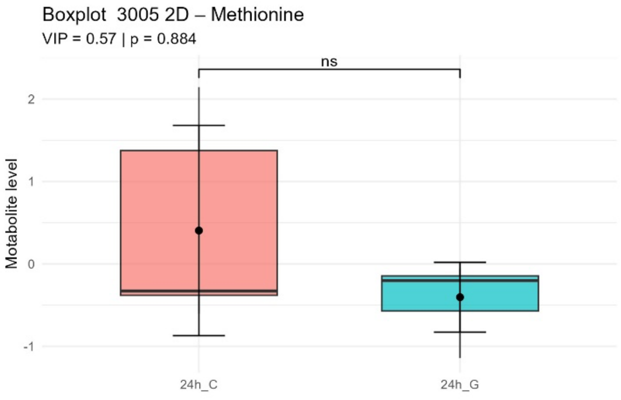
Boxplot 3005 2D – Methionine
VIP = 0.57 | p = 0.884
ns
Motabolite level
2
1
0
-1
24h_C
24h_G

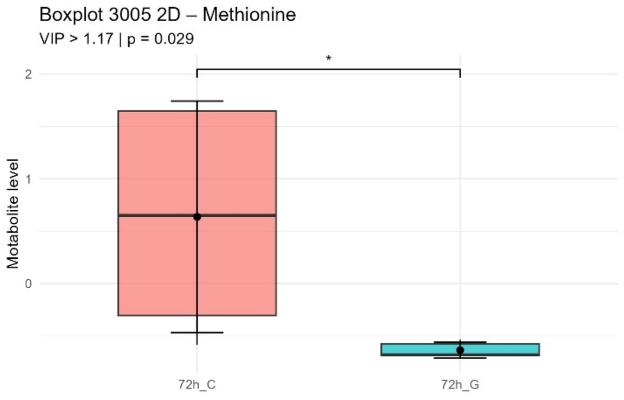
Boxplot 3005 2D – Methionine
VIP > 1.17 | p = 0.029
*
Motabolite level
2
1
0
72h_C
72h_G

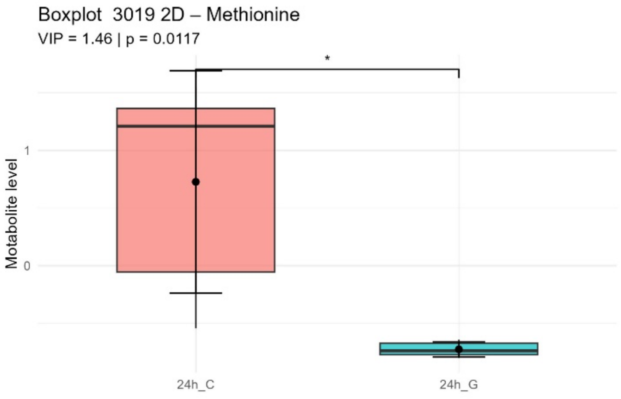
Boxplot 3019 2D – Methionine
VIP = 1.46 | p = 0.0117
*
Motabolite level
1
0
24h_C
24h_G

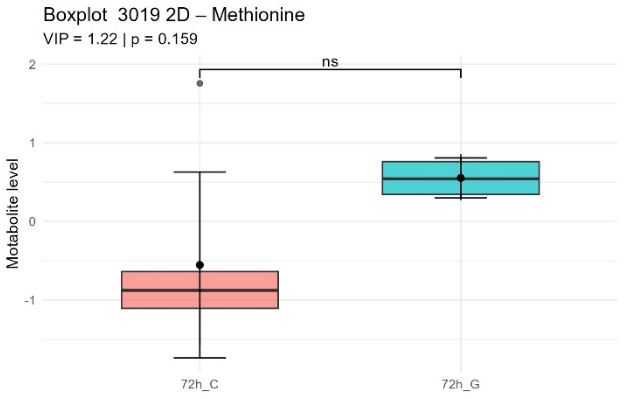
Boxplot 3019 2D – Methionine
VIP = 1.22 | p = 0.159
ns
Motabolite level
2
1
0
-1
72h_C
72h_G

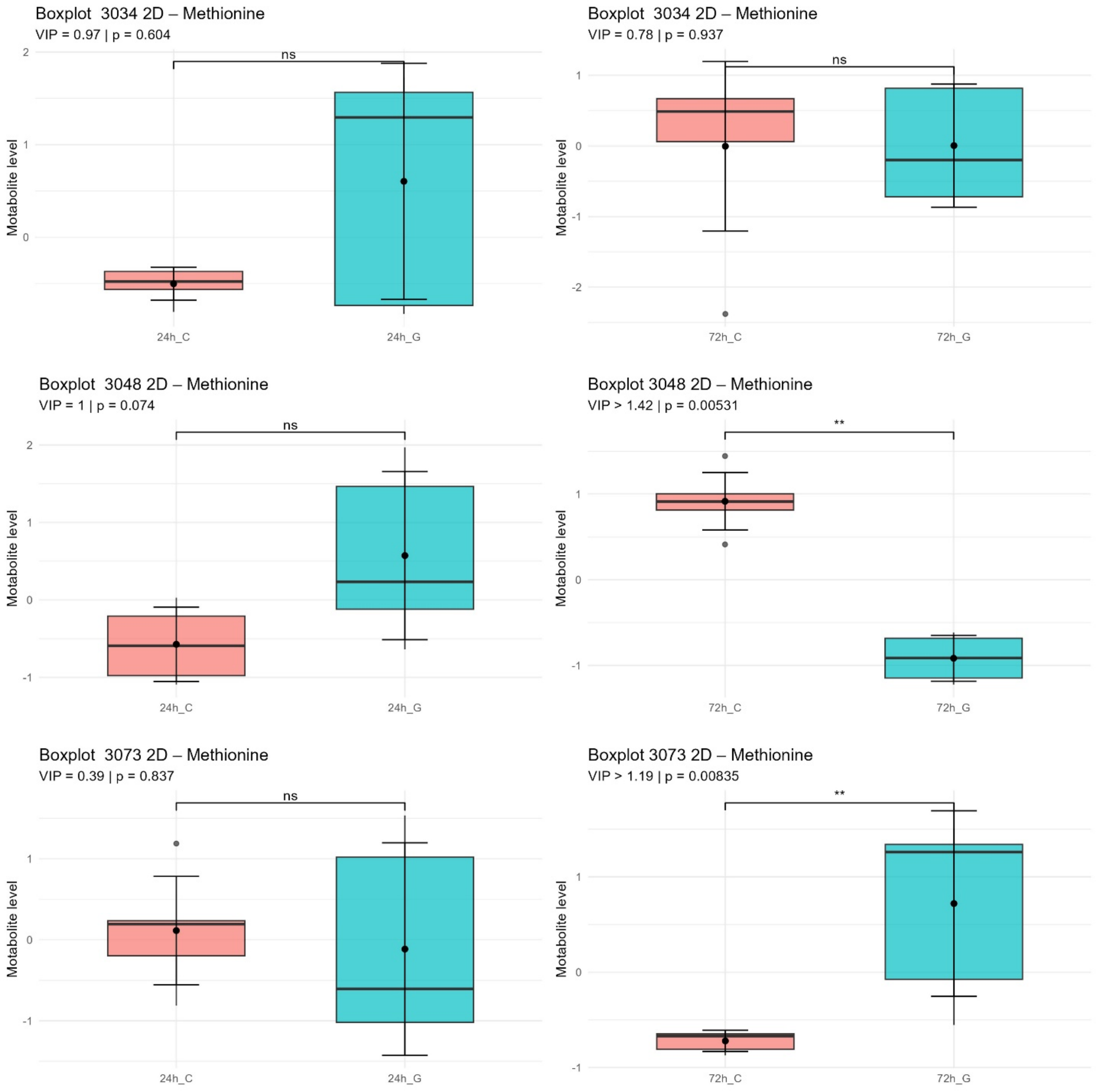

Figure S10. Change in levels of methionine between GaM treated cells and untreated control in 2D culture with VIP score and p-value (FDR).

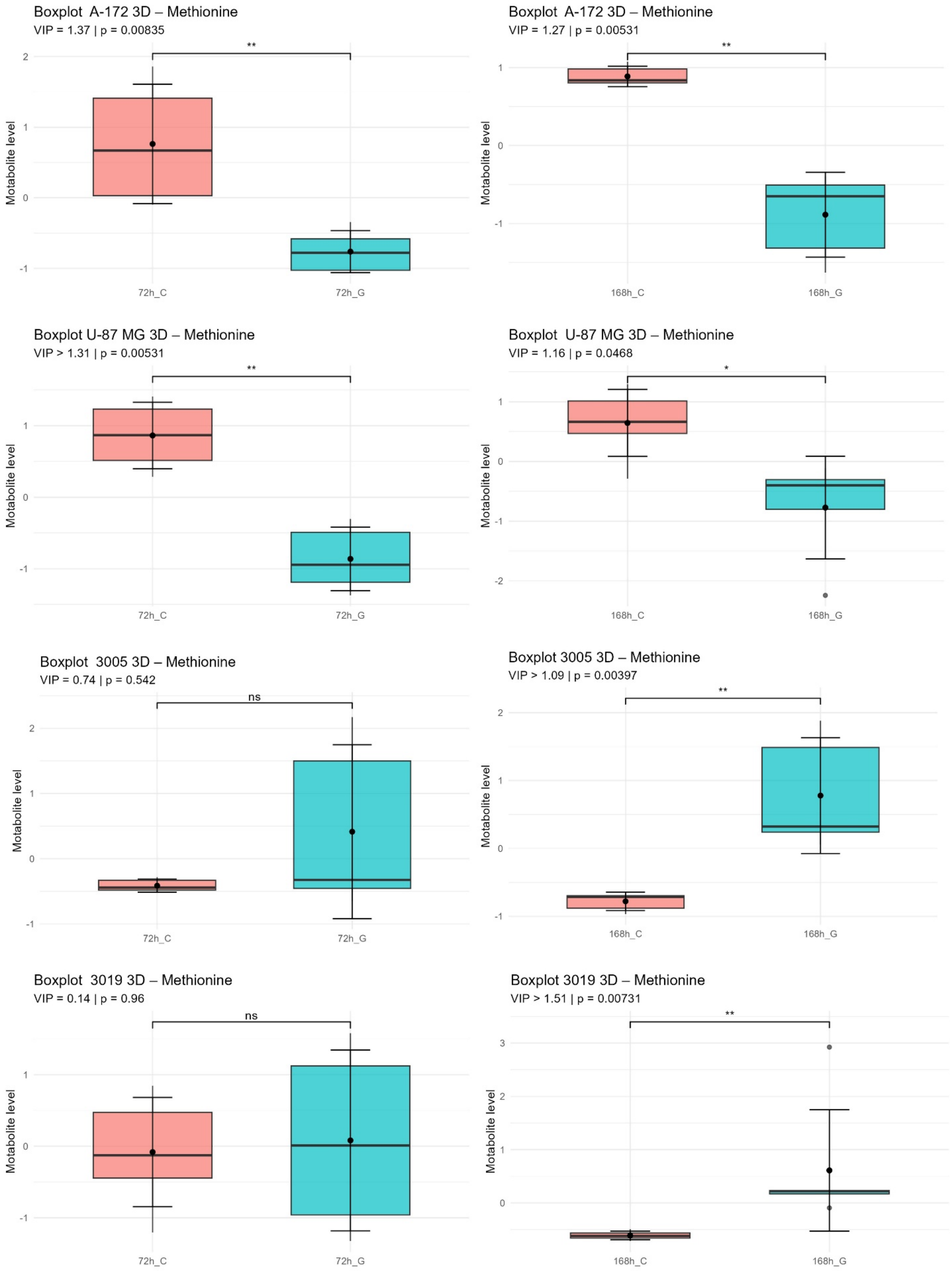
Boxplot A-172 3D – Methionine
VIP = 1.37 | p = 0.00835
**
Motabolite level
72h_C
72h_G
Boxplot A-172 3D – Methionine
VIP = 1.27 | p = 0.00531
**
Motabolite level
168h_C
168h_G
Boxplot U-87 MG 3D – Methionine
VIP > 1.31 | p = 0.00531
**
Motabolite level
72h_C
72h_G
Boxplot U-87 MG 3D – Methionine
VIP = 1.16 | p = 0.0468
*
Motabolite level
168h_C
168h_G
Boxplot 3005 3D – Methionine
VIP = 0.74 | p = 0.542
ns
Motabolite level
72h_C
72h_G
Boxplot 3005 3D – Methionine
VIP > 1.09 | p = 0.00397
**
Motabolite level
168h_C
168h_G
Boxplot 3019 3D – Methionine
VIP = 0.14 | p = 0.96
ns
Motabolite level
72h_C
72h_G
Boxplot 3019 3D – Methionine
VIP > 1.51 | p = 0.00731
**
Motabolite level
168h_C
168h_G

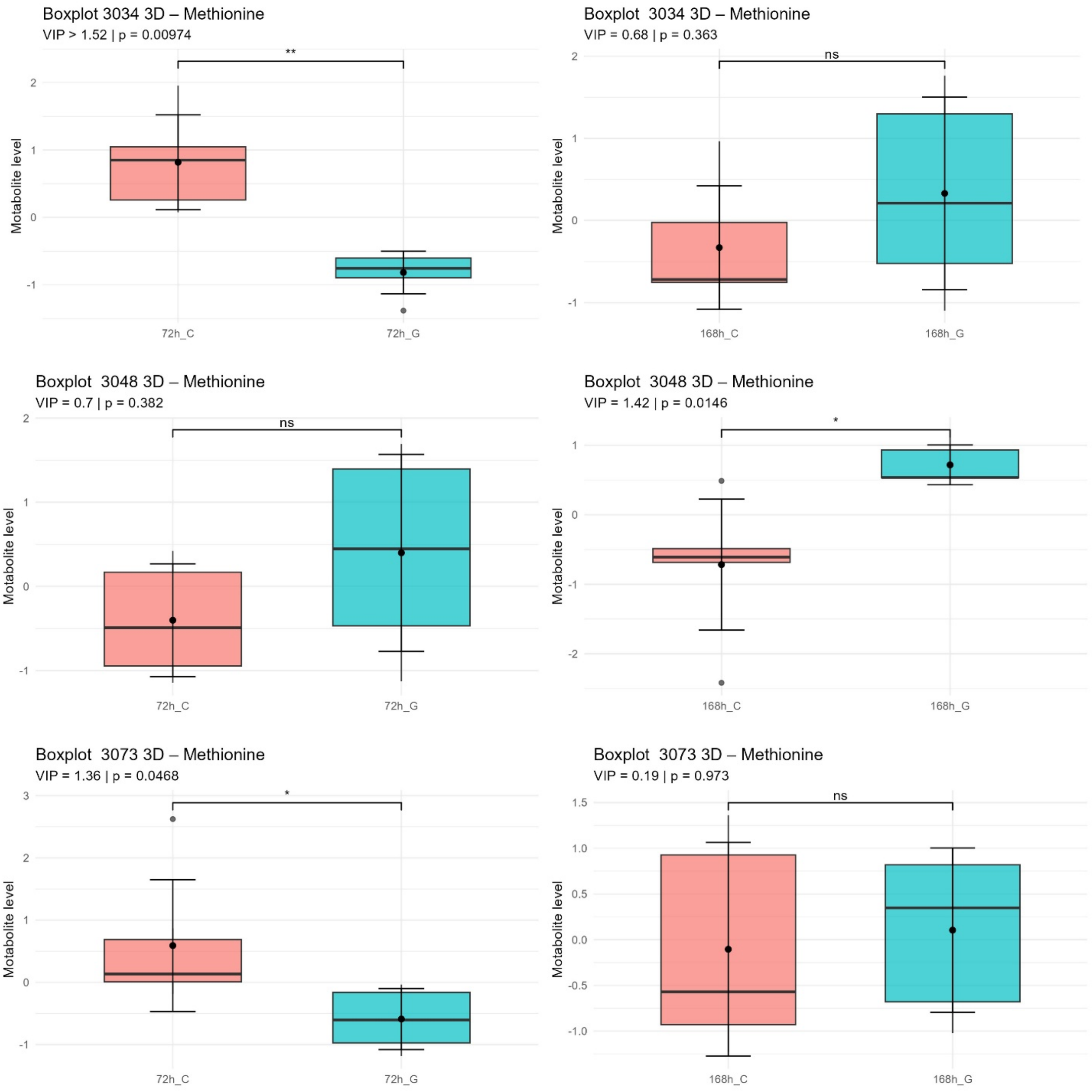

Figure S11. Change in levels of methionine between GaM treated cells and untreated control in 3D culture with VIP score and p-value (FDR).

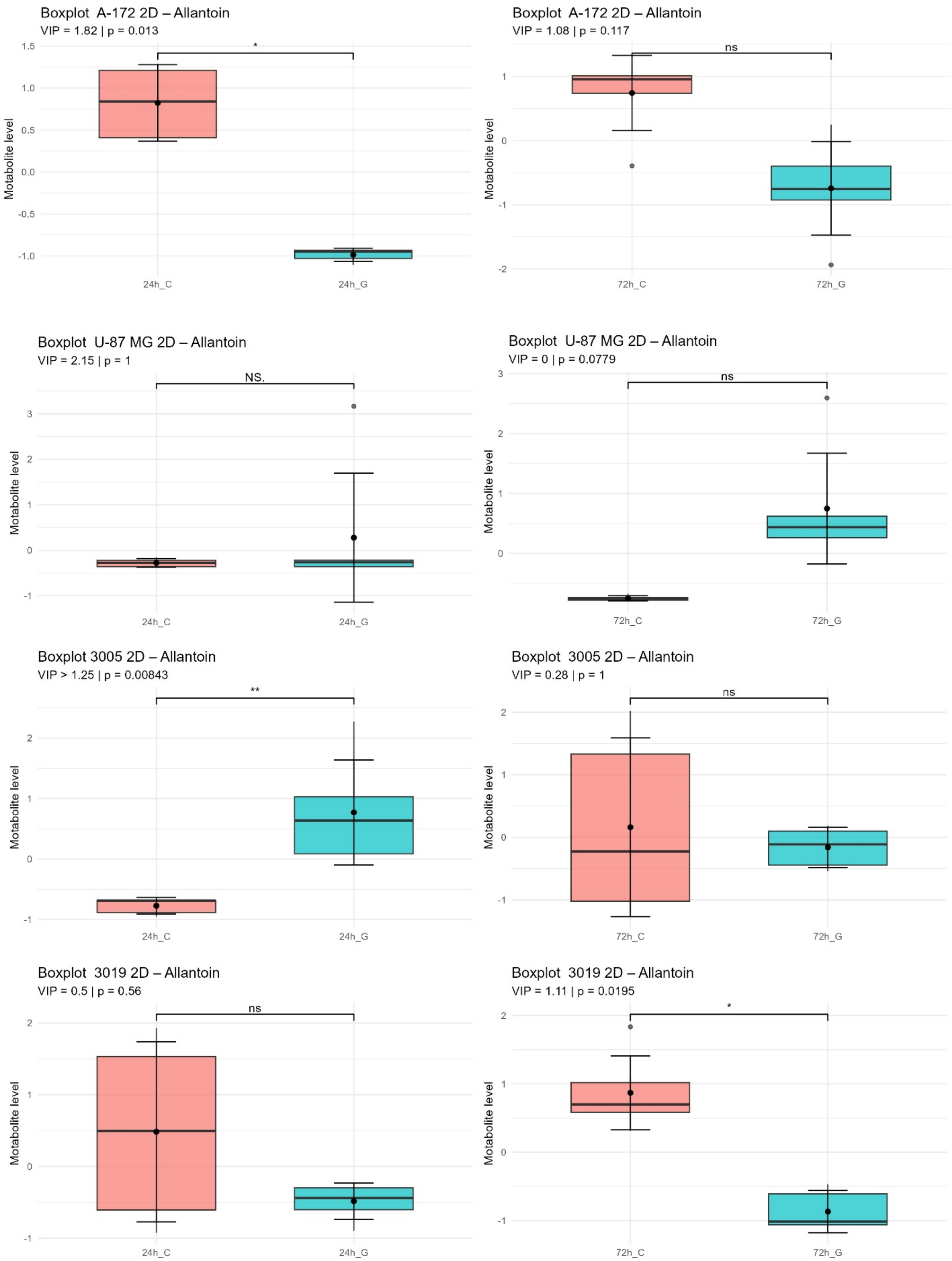
Boxplot A-172 2D – Allantoin
VIP = 1.82 | p = 0.013
*
Motabolite level
1.5
1.0
0.5
0.0
-0.5
-1.0
24h_C
24h_G
Boxplot A-172 2D – Allantoin
VIP = 1.08 | p = 0.117
ns
Motabolite level
1
0
-1
-2
72h_C
72h_G
Boxplot U-87 MG 2D – Allantoin
VIP = 2.15 | p = 1
NS.
Motabolite level
3
2
1
0
-1
24h_C
24h_G
Boxplot U-87 MG 2D – Allantoin
VIP = 0 | p = 0.0779
ns
Motabolite level
3
2
1
0
72h_C
72h_G
Boxplot 3005 2D – Allantoin
VIP > 1.25 | p = 0.00843
**
Motabolite level
2
1
0
-1
24h_C
24h_G
Boxplot 3005 2D – Allantoin
VIP = 0.28 | p = 1
ns
Motabolite level
2
1
0
-1
72h_C
72h_G
Boxplot 3019 2D – Allantoin
VIP = 0.5 | p = 0.56
ns
Motabolite level
2
1
0
-1
24h_C
24h_G
Boxplot 3019 2D – Allantoin
VIP = 1.11 | p = 0.0195
*
Motabolite level
2
1
0
-1
72h_C
72h_G

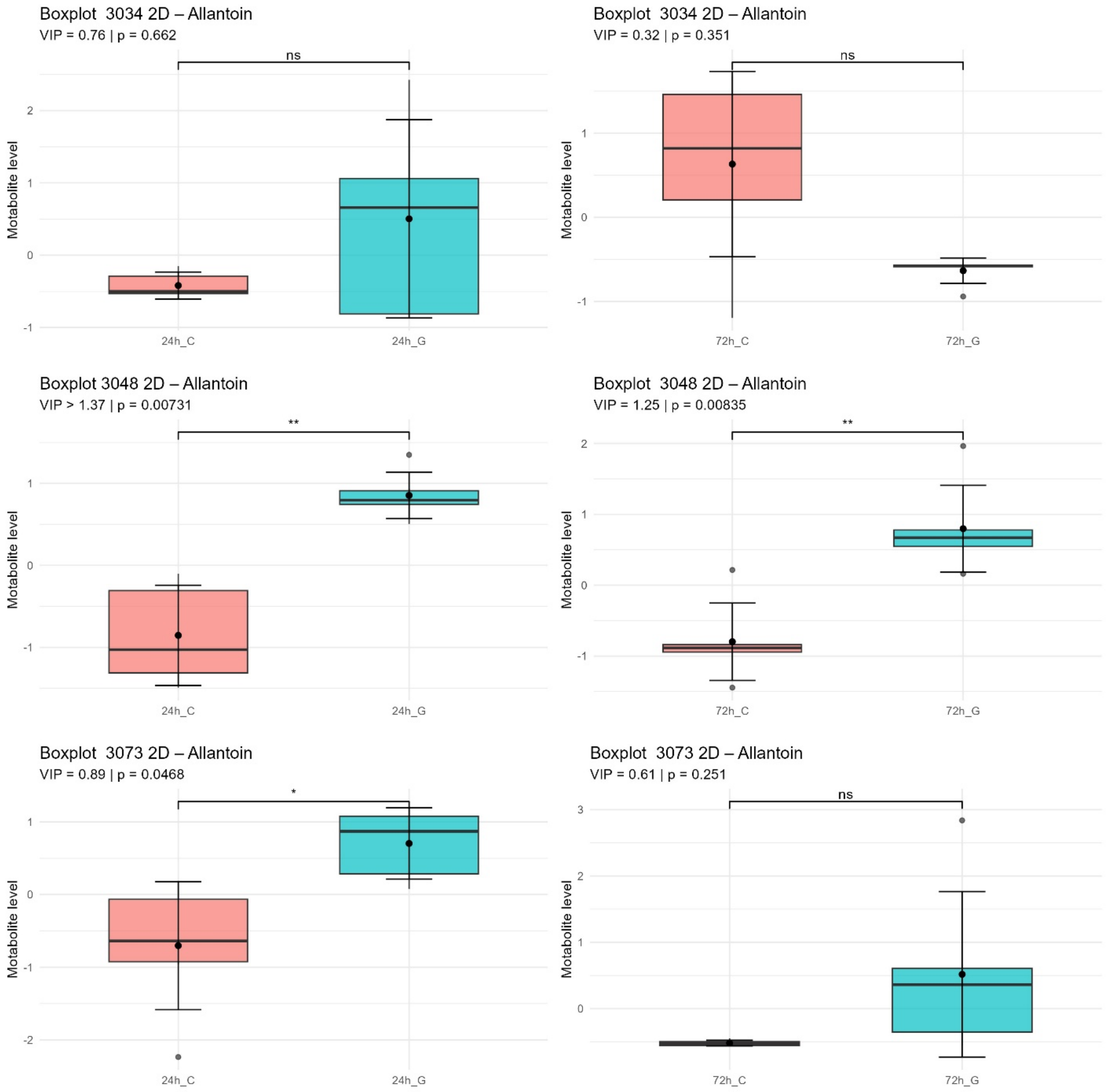

Figure S12. Change in levels of allatoin between GaM treated cells and untreated control in 2D culture with VIP score and p-value (FDR).

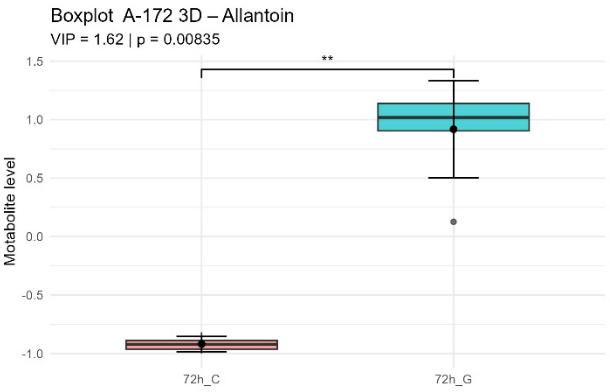
Boxplot  A-172 3D – Allantoin
VIP = 1.62 | p = 0.00835
**
Motabolite level
1.5
1.0
0.5
0.0
-0.5
-1.0
72h_C
72h_G

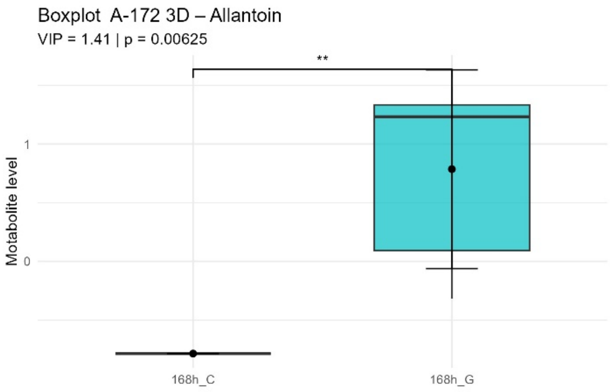
Boxplot  A-172 3D – Allantoin
VIP = 1.41 | p = 0.00625
**
Motabolite level
1
0
168h_C
168h_G

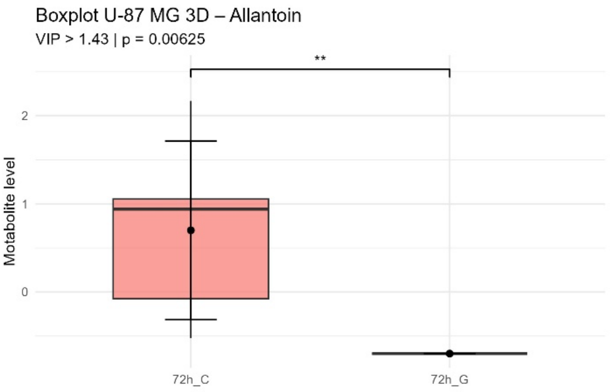
Boxplot U-87 MG 3D – Allantoin
VIP > 1.43 | p = 0.00625
**
Motabolite level
2
1
0
72h_C
72h_G

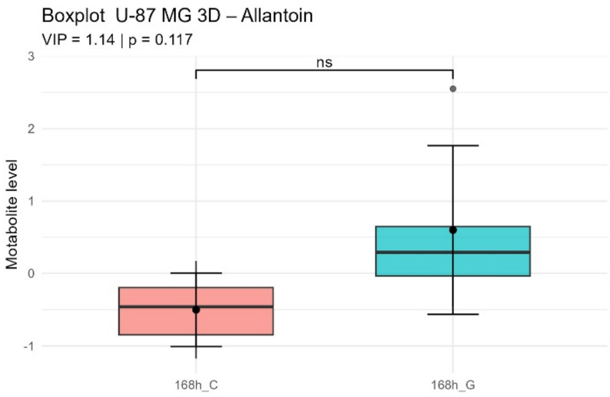
Boxplot  U-87 MG 3D – Allantoin
VIP = 1.14 | p = 0.117
ns
Motabolite level
3
2
1
0
-1
168h_C
168h_G

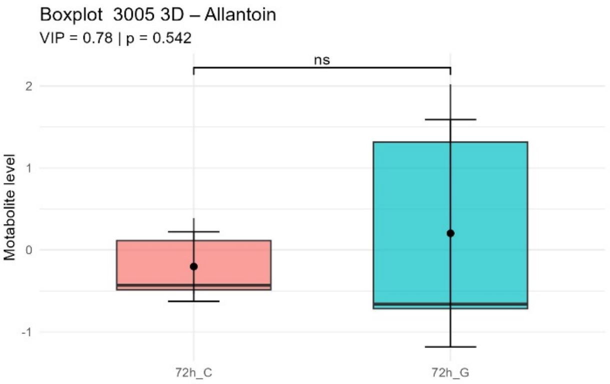
Boxplot  3005 3D – Allantoin
VIP = 0.78 | p = 0.542
ns
Motabolite level
2
1
0
-1
72h_C
72h_G

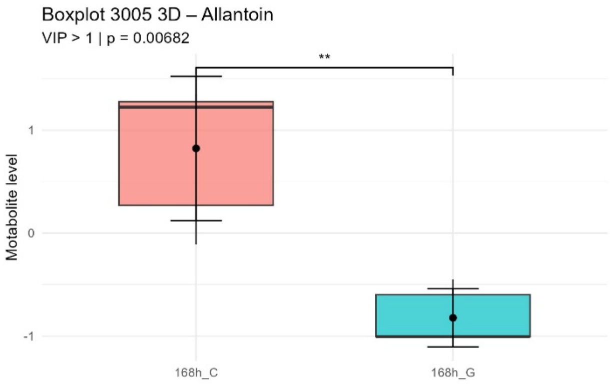
Boxplot  3005 3D – Allantoin
VIP > 1 | p = 0.00682
**
Motabolite level
1
0
-1
168h_C
168h_G

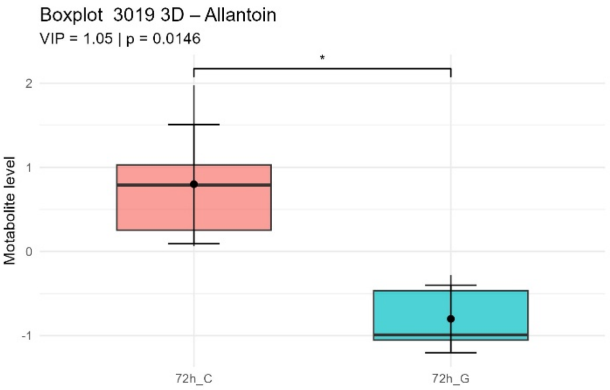
Boxplot  3019 3D – Allantoin
VIP = 1.05 | p = 0.0146
*
Motabolite level
2
1
0
-1
72h_C
72h_G

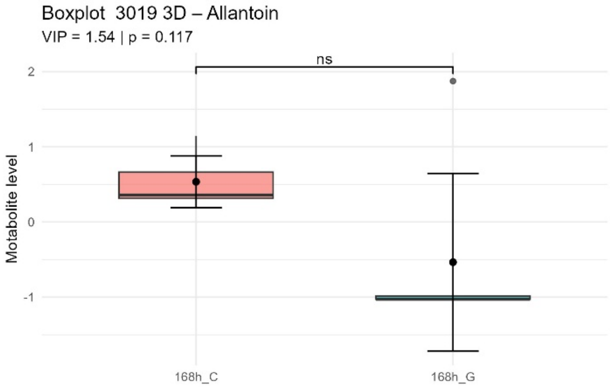
Boxplot  3019 3D – Allantoin
VIP = 1.54 | p = 0.117
ns
Motabolite level
2
1
0
-1
168h_C
168h_G

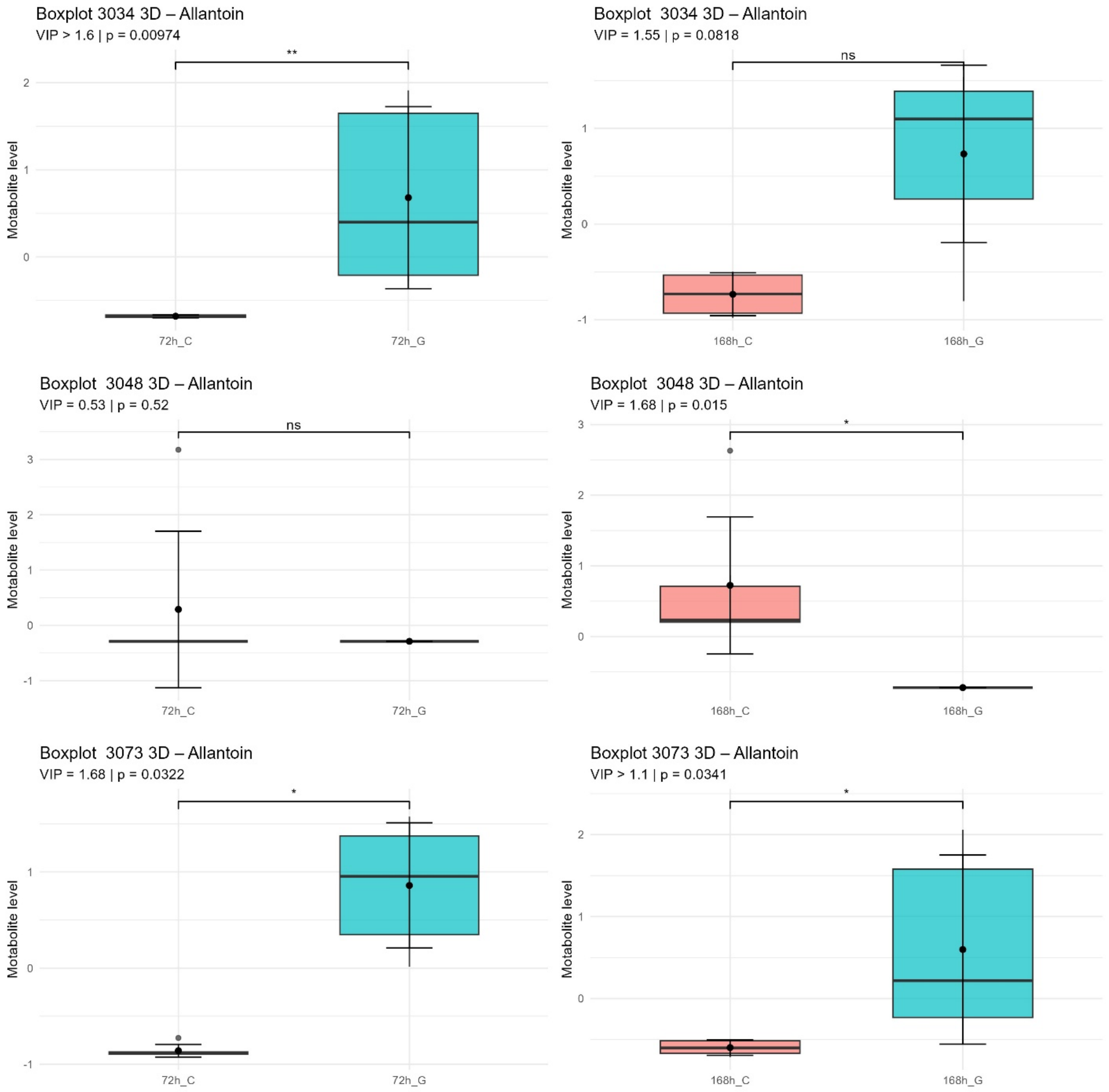

Figure S13. Change in levels of allatoin between GaM treated cells and untreated control in 3D culture with VIP score and p-value (FDR).

Code for R:

Code S1. Dose-response curve with IC90, IC50 and IC10 calculations

```
library(tidyverse)

library(drc)

library(stringr)

df_long <- df_raw %>%

  pivot_longer(cols = everything(), names_to = "Concentration", values_to = "Absorbance") %>%

  mutate(Concentration = str_remove(Concentration, "^X")) %>%

  mutate(Concentration = as.numeric(Concentration))

df_clean <- df_long %>%

  group_by(Concentration) %>%

  mutate(

    mean_abs = mean(Absorbance, na.rm = TRUE),

    sd_abs = sd(Absorbance, na.rm = TRUE),

    rel_sd = (sd_abs / mean_abs) * 100

  ) %>%

  filter(rel_sd <= 15 | abs(Absorbance - mean_abs) <= 1.5 * sd_abs) %>%

  ungroup()

df_summary <- df_clean %>%

  group_by(Concentration) %>%

  summarise(

    mean_abs = mean(Absorbance, na.rm = TRUE),

    sd_abs = sd(Absorbance, na.rm = TRUE),

    .groups = "drop"

  )

control_mean <- df_summary %>% filter(Concentration == 0) %>% pull(mean_abs)
```

```
df_summary <- df_summary %>%
  mutate(
    Survival = (mean_abs / control_mean) * 100,
    Survival_sd = (sd_abs / control_mean) * 100
  )

model <- drm(Survival ~ Concentration, data = df_summary, fct = LL.4())

ic50 <- ED(model, 50, interval = "none")
ic10 <- ED(model, 10, interval = "none")
ic90 <- ED(model, 90, interval = "none")
ic_values <- ED(model, c(10, 50, 90), interval = "none")
df_ic <- data.frame(
  Parameter = c("IC10", "IC50", "IC90"),
  Value_uM = round(ic_values, 2)
)
ggplot(df_summary, aes(x = Concentration, y = Survival)) +
  geom_point(size = 3, color = "steelblue") +
  geom_errorbar(aes(ymin = Survival - Survival_sd, ymax = Survival + Survival_sd),
              width = 2, color = "gray50") +
  stat_smooth(method = "drm", formula = y ~ x,
             method.args = list(fct = LL.4()), se = FALSE, color = "red") +
  geom_vline(xintercept = ic10[1], linetype = "dotted", color = "blue") +
  geom_vline(xintercept = ic50[1], linetype = "dashed", color = "red")  +
  geom_vline(xintercept = ic90[1], linetype = "dotted", color = "purple") +
```

```
annotate("text", x = ic10[1], y = 90, label = paste("IC10 =", round(ic10[1], 2)), color = "blue") +
annotate("text", x = ic50[1], y = 50, label = paste("IC50 =", round(ic50[1], 2)), color = "red" )  +
annotate("text", x = ic90[1], y = 10, label = paste("IC90 =", round(ic90[1], 2)), color = "purple") +
labs(
  title = "U-87MG 3D Viability (CellTiter Glo Assay)",
  x = "Concentration [µM]",
  y = "% Viability compared to control"
) +
theme_minimal()
```

Code S2. T-test for TFRC level determination and significance of changes determination in 2D and 3D control

```
library(readr)

library(tidyverse)

library(ggplot2)

library(ggpubr)

library(dplyr)

library(FSA)

cell_data <- cell_data %>% rename(Group = 1)

data_long <- cell_data %>%

  pivot_longer(cols = -Group, names_to = "CellLine", values_to = "Value") %>%

  mutate(Group = factor(Group, levels = c("C_2D", "C_3D")))

ttest_results <- data_long %>%

  group_by(CellLine) %>%

  summarise(

    p_value = t.test(Value ~ Group)$p.value

  ) %>%

  mutate(

    stars = case_when(

      p_value <= 0.001 ~ "***",

      p_value <= 0.01  ~ "**",

      p_value <= 0.05  ~ "*",
```

```
    )
  )

pval_data <- ttest_results %>%
  mutate(
    group1 = "2D",
    group2 = "3D",
    y.position = 1.1 * max(data_long$Value, na.rm = TRUE)
  ) %>%
  select(CellLine, group1, group2, y.position, stars)

ggplot(data_long, aes(x = Group, y = Value, fill = Group)) +
  stat_summary(fun = mean, geom = "bar", width = 0.7, alpha = 0.8) +
  stat_summary(fun.data = mean_sdl, geom = "errorbar", width = 0.2) +
  stat_pvalue_manual(
    pval_data,
    label = "stars",
    xmin = "group1",
    xmax = "group2",
    y.position = "y.position"
  ) +
  facet_wrap(~CellLine, scales = "free_y") +
  theme_minimal() +
  labs(title = "TFRC in non-treated cells" ,
       subtitle = "Test t-Studenta + p-value",
       x = NULL, y = "TFRC/ Total Protein") +
```

```
theme(
  axis.text.x = element_blank(),
  axis.ticks.x = element_blank(),
  legend.position = c(0.95, 0.05),
  legend.justification = c(1, 0),
  plot.title = element_text(hjust = 0.5)
)
```

Code S3. Pearson correlation test between IC10 and TFRC level in 2D and 3D cell lines.

```
library(tidyverse)
library(ggpubr)

tfrc <- df[1:3, -1] %>%
  pivot_longer(cols = everything(), names_to = "CellLine", values_to = "TFRC") %>%
  mutate(gene = "TFRC")

ic10 <- df[4:6, -1] %>%
  pivot_longer(cols = everything(), names_to = "CellLine", values_to = "IC10") %>%
  mutate(gene = "IC10")

tfrc_avg <- tfrc %>%
  group_by(CellLine) %>%
  summarise(TFRC = mean(TFRC, na.rm = TRUE))

ic10_avg <- ic10 %>%
  group_by(CellLine) %>%
  summarise(IC10 = mean(IC10, na.rm = TRUE))

tfrc_avg <- tfrc %>% group_by(CellLine) %>% summarise(TFRC = mean(TFRC, na.rm = TRUE))
ic50_avg <- ic10 %>% group_by(CellLine) %>% summarise(IC50 = mean(IC10, na.rm = TRUE))

df_corr <- left_join(tfrc_avg, ic10_avg, by = "CellLine") %>% drop_na()

# Korelacja
```

```r
cor.test(df_corr$TFRC, df_corr$IC10, method = "pearson")

ggplot(df_corr, aes(x = TFRC, y = IC10)) +
  geom_point(size = 3.2, color = "steelblue") +
  geom_smooth(method = "lm", se = FALSE, color = "darkred", linewidth = 1) +
  # jeśli chcesz klasyczny geom_text (większa czcionka):
  # geom_text(aes(label = CellLine), vjust = -0.9, size = 4.2, fontface = "italic")
  # albo lepiej – etykiety, które się nie nakładają:
  ggrepel::geom_text_repel(aes(label = CellLine), size = 4.2, fontface = "italic",
                           max.overlaps = Inf, box.padding = 0.25) +
  stat_cor(method = "pearson",
           label.x = min(df_corr$TFRC, na.rm = TRUE),
           label.y = max(df_corr$IC10,  na.rm = TRUE),
           size = 5) +  # << rozmiar napisu R i p
  labs(
    title = "3D Correlation between TFRC expression and IC10",
    x = "TFRC level (mean)",
    y = "IC10 (mean)"
  ) +
  theme_minimal(base_size = 16) +   # << bazowy rozmiar czcionki
  theme(
    plot.title  = element_text(size = 18, face = "bold"),
    axis.title  = element_text(size = 15),
    axis.text   = element_text(size = 13)
  )
```

Code S.4 TFRC level determination in treated cells and untreated control, Kurskal-Wallis test with dunn post-hoc.

```
library(readr)

library(tidyverse)

library(ggplot2)

library(ggpubr)

library(dplyr)

library(FSA)

data_long_3D <- tfrc1_3D %>%

  pivot_longer(cols = -Group, names_to = "CellLine", values_to = "TfRC_per_protein") %>%

  mutate(Group = factor(Group, levels = c("C", "72h_G", "168h_G")))

dunn_all_3D <- data_long_3D %>%

  group_by(CellLine) %>%

  group_map(~{

    kw <- kruskal.test(TfRC_per_protein ~ Group, data = .x)

    if (kw$p.value > 0.05) return(NULL)

    dunn <- dunnTest(TfRC_per_protein ~ Group, data = .x, method = "bh")$res

    dunn %>%

      separate(Comparison, into = c("group1", "group2"), sep = " - ") %>%

      mutate(

        CellLine = .y$CellLine,

        y.position = max(.x$TfRC_per_protein, na.rm = TRUE) * 1.1 + row_number() * 0.05,
```

```
      stars = case_when(
        P.adj <= 0.001 ~ "***",
        P.adj <= 0.01  ~ "**",
        P.adj <= 0.05  ~ "*",
        TRUE           ~ NA_character_
      )
    ) %>%
    drop_na(stars) %>%
    select(group1, group2, y.position, stars, CellLine)
  }) %>%
  bind_rows()

ggplot(data_long_3D, aes(x = Group, y = TfRC_per_protein, fill = Group)) +
  stat_summary(fun = mean, geom = "bar", width = 0.7, alpha = 0.8) +
  stat_summary(fun.data = mean_sdl, geom = "errorbar", width = 0.2) +
  stat_pvalue_manual(
    dunn_all_3D,
    label = "stars",
    xmin = "group1",
    xmax = "group2",
    y.position = "y.position",
    tip.length = 0.01
  ) +
  facet_wrap(~CellLine, scales = "free_y") +
  theme_minimal() +
  labs(title = "TFRC of 3D cells after treatement",
```

```
    subtitle = "Kruskal-Wallis + Dunn post-hoc (p-value)",
    x = "", y = "TFRC / Total Protein") +
theme(
  plot.title = element_text(hjust = 0.5),
  axis.text.x = element_blank(),
  axis.ticks.x = element_blank(),
  legend.position = c(0.95, 0.05),
  legend.justification = c(1, 0)
)
```

Code S5. The Oxygen Consuption Rate curve in time.

```
library(tidyverse)

names(df)[1] <- "ID"

df <- df %>%

  separate(ID, into = c("CellLine", "Condition"), sep = "_", extra = "merge")

names(df) <- gsub("^X", "", names(df))

df_long <- df %>%

  pivot_longer(

    cols = -c(CellLine, Condition),

    names_to = "Time",

    values_to = "Absorbance"

  )

df_long$Time <- as.numeric(df_long$Time)

df_summary <- df_long %>%

  group_by(CellLine, Condition, Time) %>%

  summarise(

    mean_abs = mean(Absorbance, na.rm = TRUE),

    sd_abs = sd(Absorbance, na.rm = TRUE),

    .groups = "drop"

  )
```

```
cell_to_plot <- "A-172"

df_plot <- df_summary %>% filter(CellLine == cell_to_plot)

ggplot(df_plot, aes(x = Time, y = mean_abs, color = Condition)) +
  geom_line(size = 1.2) +   # grubsza linia
  geom_point(size = 3) +    # większe kropki
  geom_errorbar(aes(ymin = mean_abs - sd_abs, ymax = mean_abs + sd_abs),
                width = 1, size = 0.6) +  # cieńsze słupki błędu
  scale_x_continuous(breaks = seq(0, 135, by = 5)) +
  labs(
    title = paste("Oxygen Consumption Rate -", cell_to_plot),
    x = "Time (min)",
    y = "Absorbancce (mean ± SD)"
  ) +
  theme_minimal()
```

Code S6 Wilcox test with FDR correction and selection of metabolites with VIP > 1 and adj. p-value (FDR).

```
names(vip)[1] <- "Metabolite"

metabolite_data <- df %>%
  select(-NAME, -Group)

log_data <- log10(metabolite_data + 1)  # dodajemy 1, by uniknąć log(0)

scaled_data <- scale(log_data)

df_scaled <- bind_cols(df %>% select(NAME, Group), as.data.frame(scaled_data))

df_long <- df_scaled %>%
  pivot_longer(-c(NAME, Group), names_to = "Metabolite", values_to = "Value")

stats <- df_long %>%
  group_by(Metabolite) %>%
  summarise(p_value = wilcox.test(Value ~ Group)$p.value) %>%
  mutate(p_adj = p.adjust(p_value, method = "fdr"))

vip_df <- vip %>%
  select(Metabolite, VIP = `Comp. 1`)

final <- stats %>%
```

```r
  inner_join(vip_df, by = "Metabolite") %>%
  filter(p_adj < 0.05 & VIP > 1) %>%
  mutate(
    stars = case_when(
      p_adj <= 0.001 ~ "***",
      p_adj <= 0.01  ~ "**",
      p_adj <= 0.05  ~ "*"
    )
  )

selected_metabolites <- final$Metabolite

all_meta <- stats %>%
  inner_join(vip_df, by = "Metabolite")

for (met in selected_metabolites) {
  df_met <- df_long %>% filter(Metabolite == met)

  if (all(c("72h_C", "72h_G") %in% unique(df_met$Group))) {

    y_max <- max(df_met$Value, na.rm = TRUE)
    p_val <- all_meta$p_adj[all_meta$Metabolite == met]

    p <- ggplot(df_met, aes(x = Group, y = Value, fill = Group)) +
      geom_boxplot(width = 0.6, alpha = 0.7) +
      stat_summary(fun = mean, geom = "point", shape = 20, size = 3, color = "black") +
```

```
    stat_summary(fun.data = mean_sdl, fun.args = list(mult = 1),
                 geom = "errorbar", width = 0.2, color = "black") +
    geom_signif(
      annotations = ifelse(p_val <= 0.001, "***",
                           ifelse(p_val <= 0.01, "**",
                                  ifelse(p_val <= 0.05, "*", "ns"))),
      y_position = y_max * 1.1,
      xmin = 1, xmax = 2
    ) +
    labs(title = paste("Boxplot  3034 3D –", met),
         subtitle = paste("VIP =", round(all_meta$VIP[all_meta$Metabolite == met], 2),
                          "| p =", signif(all_meta$p_adj[all_meta$Metabolite == met], 3)),
         y = "Motabolite level", x = "") +
    theme_minimal() +
    theme(legend.position = "none")

  ggsave(filename = paste0("boxplot_", met, ".png"), plot = p, width = 6, height = 4, dpi = 300)
 }
}
```
